# Distinct p21 dynamics drive alternative routes to whole-genome duplication through a common CDK4/6-dependent polyploid G0 state

**DOI:** 10.64898/2026.01.15.699545

**Authors:** Yian Yang, Anuraag Bukkuri, Dante Poe, Janet McLaughlin, Sihan Hao, Daniel D. Brown, Katarzyna M. Kedziora, Wayne Stallaert

## Abstract

Whole-genome duplication (WGD) fuels tumor evolution and therapy resistance, yet the molecular mechanisms governing the switch from the canonical mitotic cell cycle to the endoreplication cycle remain unclear. Here, we combine single-cell proteomics, manifold learning, and live-cell imaging to map the intersection of the mitotic and endoreplication cycles in breast cancer cells exposed to genotoxic agents. We identify two distinct routes to WGD driven by distinct p21 dynamics. High p21 induction induces G2 exit and endocycling, whereas insufficient p21 permits mitotic entry followed by slippage and endomitosis. This therapy-induced switch acts as a facultative stress response, generating drug-resistant polyploid populations that propagate genomic instability through replication stress and the generation of replication-competent micronuclei. Both paths to WGD converge on a common polyploid G0 state dependent on cyclin D1:CDK4/6 activity to complete the transition to the endoreplication cycle, revealing a shared vulnerability. Sequential treatment with genotoxic agents followed by CDK4/6 inhibitors preserves the cytotoxic efficacy of DNA-damaging drugs while simultaneously blocking entry into the endoreplication cycle and WGD-driven evolutionary rescue. These findings reveal the molecular rules governing the switch from the mitotic to endoreplication cycle and highlight the potential of WGD-blocking drugs as adjuvant therapies to inhibit drug resistance and suppress tumor evolution.

## INTRODUCTION

The endoreplication cycle is an alternative cell cycle in which cells replicate their DNA without cell division, resulting in whole-genome duplication (WGD). This process is a fundamental evolutionary mechanism across the tree of life, facilitating physiological complexity in vertebrates^1,2^, speciation and colonization of extreme environments in plants^3–5^, and adaptive stress responses in unicellular eukaryotes^6,77^. In humans, WGD plays several important physiological roles, such as facilitating liver regeneration and adaptation^8–13^, allowing for megakaryocyte differentiation and effective platelet production^14–16^, and promoting embryo implantation and maternal adaptations during pregnancy via trophoblast giant cells^17,18^. The switch from the mitotic cell division cycle to the endoreplication cycle has been co-opted for various functions across eukaryotic lineages, suggesting that conserved molecular mechanisms regulate cellular polyploidization. However, this same mechanism drives pathology. For example, polyploidization of proximal tubule epithelial cells following kidney injury promotes senescence, inflammation, and fibrosis to drive chronic kidney disease^19–22^. Similarly, WGD in cardiomyocytes, though critical for postnatal maturation, can compromise cardiac regenerative ability in adults^23–26^.

In cancer, this alternative cell cycle has been co-opted to promote oncogenesis and therapy resistance. WGD is observed across multiple solid cancers, including breast cancer, and is associated with increased mutation burden, chromosomal instability, copy number variation, drug resistance, and poor prognosis^27–29^. Furthermore, anticancer treatment with genotoxic drugs can itself promote WGD^30–35^. By buffering lethal DNA damage and accelerating adaptation due to increased genomic content, WGD provides a pernicious path to evolutionary rescue by allowing cancer cell populations to persist and adapt to DNA-damaging therapies^33,36,37^. Despite the significant impacts of WGD on cancer progression and treatment outcomes, the mechanisms underlying the switch from the canonical mitotic cell cycle to the endoreplication cycle remain unclear.

Here, we investigate the molecular mechanisms that govern the switch from the mitotic to the endoreplication cycle in breast cancer cells treated with genotoxic agents, including PARP1/2 inhibitors and the topoisomerase II inhibitor doxorubicin. Using single-cell proteomics, cell cycle mapping, and live-cell imaging, we identify two distinct routes to WGD, differentiated by distinct p21 dynamics, that ultimately converge on a common CDK4/6-dependent polyploid G0 state. This switch produces a drug-resistant cellular state that fuels tumor evolution through multiple mechanisms of genomic instability, including progressive increases in ploidy, increased replication stress, and the generation of asynchronous replication-competent micronuclei. Finally, we show that sequential therapy with CDK4/6 inhibitors preserves the cytotoxic effect of genotoxic therapies while blocking endoreplication, thereby reducing the genomic instability that typically accompanies their use.

## RESULTS

### PARP inhibition induces WGD in breast cancer

PARP1/2 inhibitors (PARPi) induce the accumulation of DNA damage by inhibiting single-strand break repair and increasing replication stress due to PARP retention^38–42^. Recently, PARPi have been shown to drive WGD in ovarian cancers^30^. We found that treatment with IC_80_ concentrations (**Extended Data Fig. 1a-b**) of the PARPi talazoparib for 5 days also induced significant increases in ploidy across a panel of breast cancer cell lines of diverse molecular and histological subtypes (**Fig. 1a**). After talazoparib treatment, the proportion of polyploid cells ranged from 25% in BCK4 cells to 65% in MDA-MB-453 (MB453) cells, with increases in polyploidy as high as 20- and 27-fold in T47D and MCF7 cell lines, respectively (**Extended Data Fig. 1c**). In both normal and cancer cells, WGD following DNA damage is often associated with cell cycle exit from G2 and subsequent endocycling^43–46^. Using immunofluorescence (IF) measurements of DNA content and retinoblastoma (RB) phosphorylation (pRB, as a marker of actively proliferating cells), we quantified the proportion of cell cycle exit (pRB-negative cells) from G1 (DNA content = 2N) versus G2 (DNA content = 4N). Talazoparib induced G2 exit in all cell lines, ranging from 21.5% of arrested BCK4 cells to 67.1% of arrested MB453 cells (**Fig. 1b**). Furthermore, ploidy progressively increased during treatment (**Fig. 1c-d, Extended Data Fig. 1d-e**) in all cell lines (**Fig. 1e, Extended Data Fig. 1f**), indicating that the mechanisms of talazoparib-induced WGD remain active during sustained PARPi treatment.

**Fig. 1:**
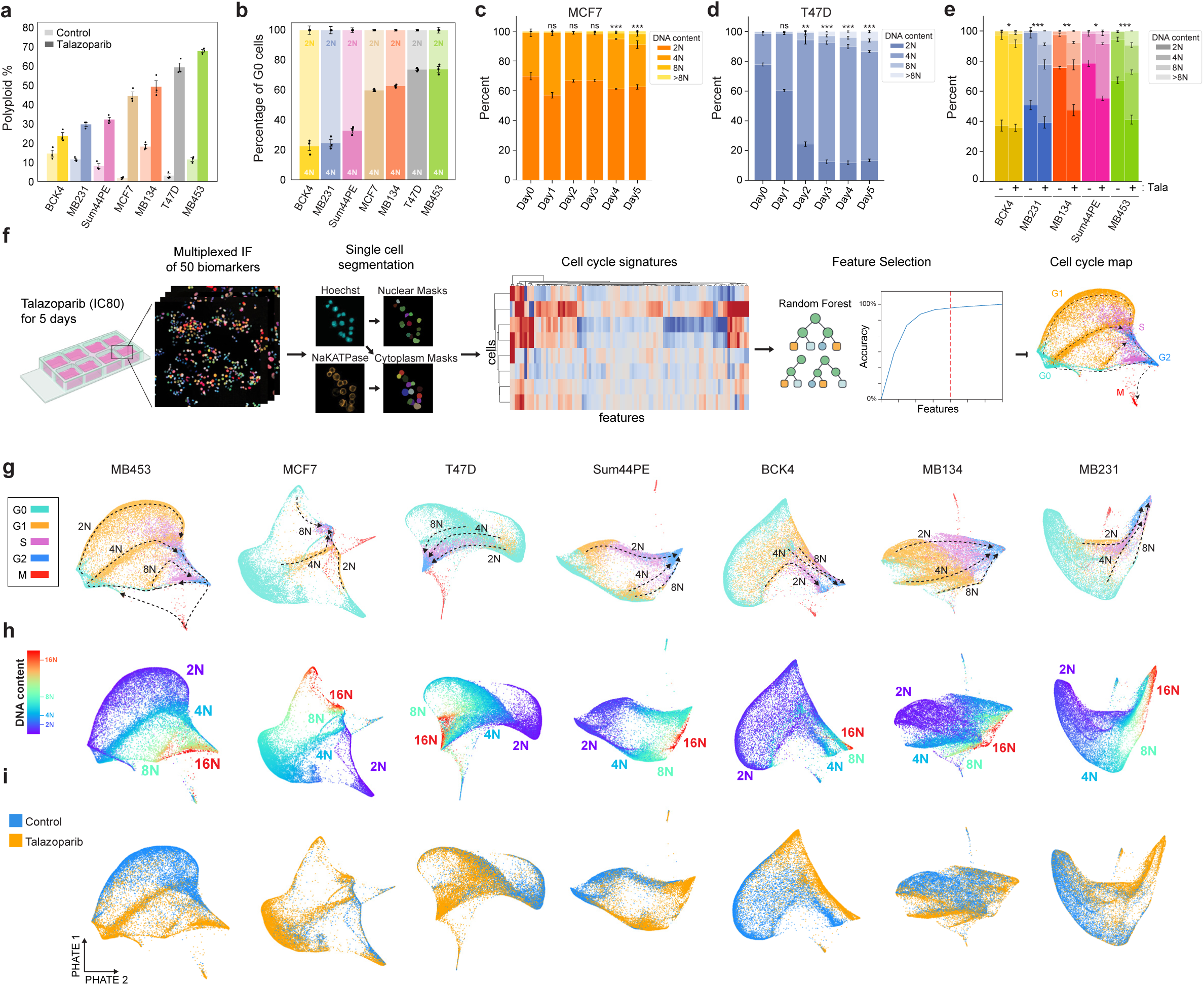
PARP1/2 inhibition induces WGD in a diversity of breast cancer models. a, Quantification of polyploid cells by single-cell IF (see *Methods*) in control and talazoparib-treated cell lines (IC_80_ concentrations for 5 d). **b**, Quantification of G1 and G2 arrest by single-cell IF following talazoparib treatment based on DNA content (2N: light bars vs 4N: dark bars) in pRB-negative arrested cells. **c-e**, DNA content by single-cell IF following talazoparib treatment: time course in MCF7 (**c**), time course in T47D cells (**d**) and after 5 days of treatment in other cell lines (**e**). **f**, Schematic of multiplexed IF and cell cycle mapping pipeline. **g-i**, Cell cycle maps of breast cancer cell lines. Single cells colored by cell cycle phase (**g**), DNA content (**h**), and treatment labels (**i**). Cell line abbreviations: MDA-MB-453 (MB453), MDA-MB-134 (MB134), MDA-MB-231 (MB231). In **a-e**, data represent mean ± s.d. from three biological replicates. In **c-e**: statistical significance (*: p < 0.05, **: p < 0.01, ***: p < 0.001) was determined using a one-way ANOVA with Tukey’s *post hoc* test comparing Days 1-5 to day 0 (**c-d**) and a two-tailed Student’s *t*-test (**e**). Talazoparib IC_80_ concentrations, MCF7: 300 nM; T47D: 500 nM; BCK4: 1 μM; Sum44PE: 428 nM; MB134: 1 μM; MB453: 1 μM; MB231: 1 μM

To identify the molecular mechanisms that control PARPi-induced WGD, we performed cell cycle mapping (**Fig. 1f**). This approach combines highly multiplexed immunofluorescence (mIF) single-cell imaging with manifold learning to visualize the cell cycle^47^ and its response to stress^48,49^. Briefly, each cell line was cultured with IC_80_ concentrations of talazoparib for 5 days, followed by chemical fixation and iterative staining and imaging of 50 core effectors of cell cycle regulation and the DNA damage response. To aid cell adhesion, we developed a novel method of sample preparation for mIF based on the principles of formalin fixation and paraffin embedding (FFPE) of tissue specimens. This “*in situ* paraffin embedding” step proved crucial in preventing cell loss when performing mIF on cell line samples (see **Methods**). For each cell in these mIF images, we quantified the nuclear and cytoplasmic abundance of each protein, DNA content, and nuclear and cytoplasmic area. This resulted in 103 single-cell proteomic measurements in >500,000 total cells that describe the response of 7 breast cancer cell lines to PARP inhibition (**Extended Data Fig. 2**). Using these proteomic signatures, cell cycle phase labels were assigned to each cell using a sequential gating strategy of conserved markers for each cell cycle phase (G0/G1/S/G2/M) (**Extended Data Fig. 1g**). We employed a random forest-based feature selection approach^47,48^ to enrich for single-cell measurements that vary in a cell cycle-dependent manner and performed manifold learning using Potential of Heat-diffusion for Affinity-based Trajectory Embedding (PHATE)^50^ to project this high-dimensional representation of cell cycle progression onto a 2-dimensional surface. These cell cycle maps visualize the diversity of paths cells take through the cell cycle within a given population (**Fig. 1g-i**). Overlaying cell cycle phase labels (**Fig. 1g**) and DNA content measurements (**Fig. 1h**) onto these maps, we resolved distinct trajectories through the cell cycle for both diploid and higher ploidy cells in all cell lines and observed an enrichment of talazoparib-treated cells along the higher ploidy trajectories (**Fig. 1i**). The map for MB453 cells, which have the highest proportion of polyploid cells following talazoparib treatment (**Fig. 1b**), most clearly demonstrates the progressive endoreplication cycles (rounds of DNA replication without cell division) that generate higher levels of ploidy (≥8N) in the sustained presence of talazoparib (**Fig. 1g**).

### PARPi-induced WGD occurs via endocycling and endomitosis

WGD due to endoreplication can occur via multiple mechanisms, producing either mononucleated or binucleated polyploid cells^51^. Whether cells become mononucleated or binucleated depends on when they exit the mitotic cell cycle (**Fig. 2a**). If cells exit during G2 (endocycling) or the early stages of mitosis (mitotic slippage or early endomitosis), a single nucleus reforms, resulting in a mononucleated cell. In contrast, exit during the later stages of mitosis (late endomitosis) or during cytokinesis (acytokinetic mitosis) leads to the formation of a single binucleated cell^51^. In response to PARPi, we primarily observed the formation of large, mononucleated polyploid cells across all cell lines (**Fig. 2b-c, Extended Data Fig. 1h-m**). To directly monitor cell cycle progression and determine the mechanisms of WGD during sustained talazoparib treatment, we performed time-lapse imaging of MCF7 and T47D cells stably expressing three fluorescent cell cycle biosensors: proliferating cell nuclear antigen (PCNA-mTq2) to annotate transitions among proliferative cell cycle phases (G1/S/G2)^52^ and kinase translocation reporters measuring CDK4^53^ and CDK2^54^ activity to identify cell cycle exit and re-entry to/from G0 based on the loss and subsequent resumption of CDK activity, respectively (**Fig. 2d**). Fully automated single-cell tracking and annotation of atypical cell cycle progression is a challenge for current methods and often requires extensive manual verification. We developed TrackGardener, a human-in-the-loop single-cell tracking analysis platform that simultaneously visualizes time-lapse images, biosensor quantification, and cell lineages for individual cell tracks, streamlining manual verification (see **Methods**). For both MCF7 (**Fig. 2e**) and T47D (**Fig. 2f**), ≥100 individual cells were tracked in untreated and talazoparib-treated conditions starting from their birth from cell division and ending either at their own cell division in mitosis or at the end of the 5 d experiment in cells that do not complete mitosis. For each single-cell track (each row in **Fig. 2e-f**), cell cycle phase transitions were annotated and plotted. In MCF7, very few cells successfully completed mitotic cell division during 5 d of talazoparib treatment and the average cell cycle duration in these cells was prolonged compared to untreated cells, primarily due to an extended G0 phase following cell division (**Fig. 2e**). Instead, 20% of talazoparib-treated MCF7 cells exited the mitotic cell cycle after cell division into a diploid G0 arrest state and 1% of cells underwent apoptosis. In 41% of talazoparib-treated MCF7 cells, we observed an abrupt reduction in CDK2 activity during G2 (**Fig. 2g-h**), resulting in mitotic cell cycle exit to a polyploid G0 state (**Fig. 2e**). Over 5 d of tracking, 20% of these cells successfully re-entered the cell cycle from polyploid G0 after highly variable durations (accompanied by an increase in CDK activity) and proceeded into a second S phase (as indicated by PCNA foci), demonstrating that PARP-induced WGD in MCF7 cells occurs primarily via endocycling (**Movie S1**). While many of these cells remain in polyploid G0 state after 5 d of observation, we classify these as endocycling based on the timing of cell cycle exit during G2 and their capacity to continue through to polyploid G1/S with prolonged observation. In T47D, we did not observe any cells successfully completing cell division through mitosis during talazoparib treatment (**Fig. 2f**). Talazoparib induced prolonged G2 durations in nearly all T47D cells (**Fig. 2f,i-j, Movie S2**), with 34% of cells remaining in a G2 arrest state after 5 d of treatment. However, 30% of T47D cells exited the mitotic cycle to polyploid G0 during these extended G2 phases and proceeded to WGD by endocycling (**Fig. 2f,k-l, Movie S3**). On the other hand, some cells successfully progressed from a stalled G2 into mitosis (as indicated by nuclear envelope breakdown and cell rounding) (**Fig. 2f,m-n, Movie S4**). These cells remained in a prolonged mitotic state (10-35 h compared to <1 h in untreated cells), exhibiting a range of defects in spindle formation and chromosomal organization (**Extended Data Fig. 3**), before reforming a single nucleus and reverting to an interphase-like cell morphology, consistent with mitotic slippage^51^, and low CDK activity indicating cell cycle arrest. From this polyploid G0 state, several cells re-entered the cell cycle in G1 and proceeded to a second S phase without cell division, thus revealing endomitosis as an additional source of WGD in T47D cells. Of note, we expect that many of the T47D cells remaining in a state of G2 arrest after 5 d would also continue on to endocycling or endomitosis with prolonged observation.

**Fig. 2:**
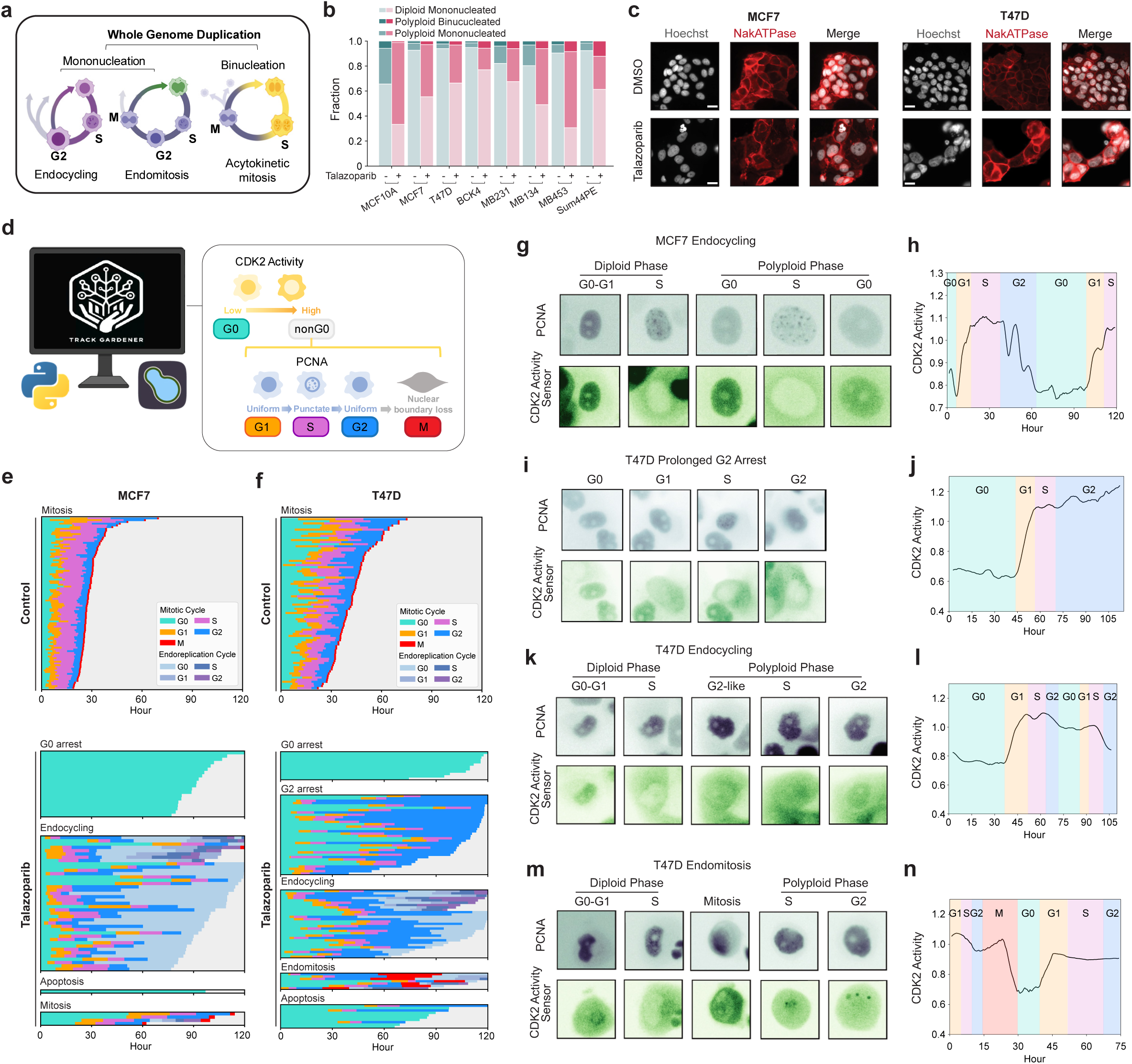
PARP1/2 inhibition induces WGD by endocycling and endomitosis in breast cancer cells. **a**, Mechanisms of WGD by endoreplication. **b-c**, Quantification of ploidy and nuclear status of control and talazoparib-treated cells (IC_80_ concentrations, 5 d) from single-cell IF (**b**) and example images; scale bar: 20 μM (**c**). **d**, Schematic of analysis and annotation strategy for single-cell tracking from time-lapse imaging. **e-f**, Time traces for MCF7 (**e**) and T47D cells (**f**) obtained by time-lapse imaging for control and talazoparib-treated conditions. Each row represents a single-cell trace colored by cell cycle phase. Cells were manually annotated by cell fate (see *Methods*). (**g-n**) Example images of PCNA-mTq2 and DHB-mVenus (CDK2 activity sensor) (**g,i,k,m**) and time traces of CDK2 activity with cell cycle phase annotations (**h,j,i,n**) of cells undergoing endocycling (**g-h, k-l**), G2 arrest (**i-j**), and endomitosis (**m-n**). Talazoparib concentrations, MCF7: 300 nM; T47D: 500 nM.

### Endocycling and endomitosis converge on a single, arrested polyploid G0 state

To extend our findings to other classes of genotoxic therapies, we investigated the mechanisms of WGD in response to treatment with doxorubicin, an FDA-approved chemotherapeutic agent for breast cancer that induces DNA damage via disruption of topoisomerase II-mediated DNA repair and the generation of reactive oxygen species^55,56^. Doxorubicin induced G2 exit (**Fig. 3a**) and WGD (**Fig. 3b-e, Extended Data Fig. 4a-b**) to a greater extent than talazoparib (**Fig. 1a-b**) in all cell lines. Both talazoparib and doxorubicin induced a concentration (**Extended Data Fig. 4d-g**) and time-dependent (**Extended Data Fig. 4h-k**) accumulation of polyploid cells, with the majority appearing after 4-5 days of treatment.

**Fig. 3:**
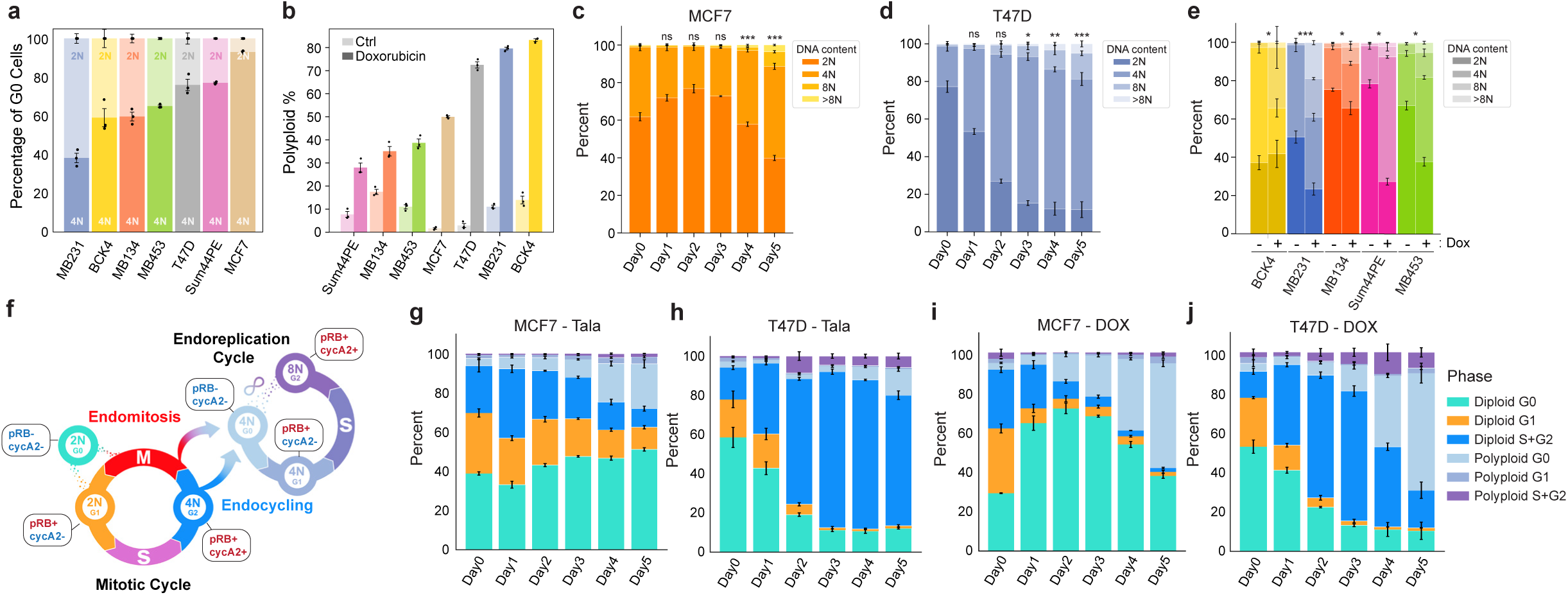
Doxorubicin and talazoparib promote a time-dependent switch to the endoreplication cycle. **a**, Quantification of G1 and G2 arrest following doxorubicin treatment based on DNA content (2N vs 4N) in pRB-negative arrested cells. **b**, Quantification of polyploid cells by single-cell IF measurements in control and doxorubicin-treated (5d) cell lines. **q-s**, Time-course of DNA content (Hoeschst) measurements by IF in MCF7 (**q**) and T47D cells (**r**) and after 5 days of doxorubicin treatment in other cell lines (**s**). **t**, Schematic of cell state transitions in the mitotic and endoreplication cycle and molecular signatures to identify each state. **u-x**, Time course of cell state transitions in MCF7 and T47D cells after talazoparib (**u-v**) or doxorubicin treatment (**w-x**). In **o-s** and **u-x**, data represent mean ± s.d. from three biological replicates. Statistical significance (*: p < 0.05, **: p < 0.01, ***: p < 0.001) was determined using a one-way ANOVA with Tukey’s *post hoc* test comparing Days 1-5 to day 0 (**c-d**) and a two-tailed Student’s *t*-test (**e**). Talazoparib concentrations, MCF7: 300 nM; T47D: 500 nM. Doxorubicin concentrations, MCF7: 100 nM; T47D: 120 nM.

Using IF measurements of pRB, cyclin A2, and DNA content, we quantified the temporal progression of cells through key cell cycle states during the transition from the mitotic cycle to the endoreplication cycle (**Fig. 3f, Extended Data Fig. 4l**). In MCF7 cells, talazoparib induced a gradual accumulation of polyploid cells in the endoreplication cycle and a concomitant loss of proliferative diploid G1 and S/G2 cells over 5 days of treatment, consistent with WGD primarily by endocycling (**Fig. 3g**). T47D cells accumulated in diploid S/G2 following talazoparib, consistent with a slower transition from the mitotic cycle to the endoreplication cycle (**Fig. 3h**). Doxorubicin showed similar trends in both cell lines, including an enrichment of S/G2 cells in T47D cells (**Fig. 3i-j**), but stimulated a faster accumulation of polyploid cells compared to talazoparib. This suggests that the mechanism that determines the path to WGD is common to both PARP inhibitors and genotoxic chemotherapy but occurs with faster kinetics following doxorubicin treatment.

To understand the molecular mechanisms that determine these paths to WGD, we performed trajectory inference on single-cell proteomic signatures obtained by mIF following talazoparib and doxorubicin treatment. We subsampled cells with 4N DNA content and annotated G2 (cycB1+/pRB+/pH3-), M (pH3+), and G0 (pRB-/pH3-) cells, generating cell cycle maps focused on the fate trajectories of G2 cells (**Fig. 4**). In both MCF7 and T47D cells, diffusion pseudotime calculations revealed two distinct trajectories emanating from G2: one leading directly into G0 (pRB-) consistent with endocycling, and another that transits through mitosis (pH3+) before exiting to G0, consistent with mitotic slippage/endomitosis^57,58^ (**Fig. 4a-d,f-i**). Using Leiden clustering, we detected distinct molecular signatures along each route and quantified the relative proportions of WGD due to endocycling versus endomitosis across our cell lines (**Fig. 4e,j, Extended Data Fig. 5a-e**). Cells transiting along the endocycling trajectory into G0 exhibited elevated p21 expression (**Fig. 4l-m**), a CDK inhibitor previously shown to promote endocycling in response to replicative stress^48,59^, whereas endomitosis was distinguished by a decrease in cyclin B1 expression (**Fig. 4n-o**), consistent with mitotic slippage^57^. Given the inherent temporal ordering of cell cycle exit from G2 versus M phase, these observations support a model in which the choice of WGD route is determined by the capacity of a cell to generate a p21 induction during G2 in response to DNA damage: Cells that successfully upregulate p21 are directed to WGD by endocycling, whereas cells that do not sufficiently upregulate p21 proceed to WGD through endomitosis (**Fig. 4k**). Supporting this model, doxorubicin, which induced a stronger p21 response than talazoparib in both cell lines (**Fig. 4p-q**), promoted faster G2 exit (**Fig. 3i-j**) and biased cells toward endocycling in both MCF7 and T47D cells (**Fig. 4r-s, left and middle panels**). Conversely, siRNA-mediated knockdown of p21 inhibited G2 exit (**Fig. 4t-u**) and biased cells towards endomitosis (**Fig. 4r-s, right panels**). The induction of p21 in response to DNA damage typically occurs in a p53-dependent manner^60^. MCF7 cells, which express wildtype p53, generated the largest p21 responses and exhibited the strongest bias towards endocycling. The remaining cell lines, which possess diverse missense/truncating p53 mutations^61^, elicited comparatively weak p21 responses (**Extended Data Fig. 5f**), exhibited longer G2 durations in response to genotoxic agents (**Fig. 2f, Extended Data Fig. 5g**) and (with the exception of MDA-MB-231 cells) showed a stronger bias towards endomitosis than MCF7 cells (**Extended Data Fig. 5a-e**). Notably, both endocycling and endomitosis occurred in all cell lines, indicating that the route to WGD is not exclusively determined by p53 status. For example, in T47D cells, which generated a weak but significant p21 response (**Fig. 4q**) despite carrying a p53 missense mutation, we observed both WGD routes and a clear p21 induction in the subpopulation of cells that transit along the endocycling trajectory (**Fig. 4m**).

**Fig. 4:**
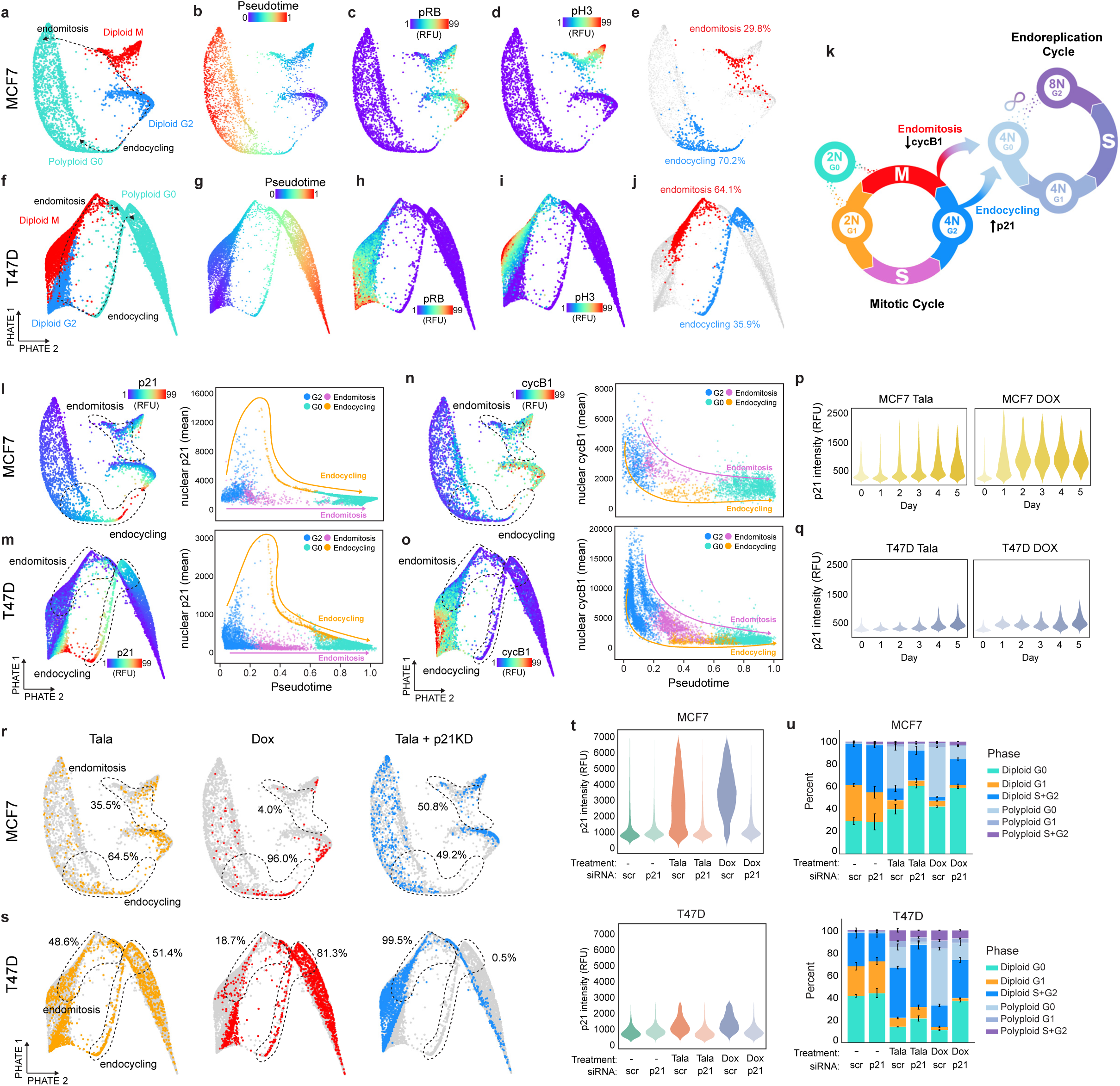
Decision between endocycling and endomitosis is governed by p21 dynamics. **a-j**, Cell cycle maps of MCF7 and T47D cells showing bifurcation of diploid G2 cell into diploid M and polyploid G0 (**a,f**), pseudotime (**b,g**), nuclear phospho-RB (**c,h**), nuclear phospho-H3 (**d,i**), and quantification of cells transiting along the endocycling and endomitosis trajectories (**e,j**). **k,** Schematic of cell state transitions in the mitotic and endoreplication cycle and the mechanisms governing endocycling and endomitosis. **l-o**, Cell cycle maps (**left panels**) overlaid with nuclear p21 (**l,n**) and nuclear cyclin B1 intensities (**m,o**) and quantification of nuclear p21 and cyclin B1 along endocycling and endomitosis trajectories (**right panels**). **p-q**, Time-course of nuclear p21 intensity by single-cell IF following talazoparib (left panels) or doxorubicin (right panels) in MCF7 (**p**) and T47D cells (**q**). **r-s**, Cell cycle maps of MCF7 (**r**) and T47D cells (**s**) quantifying the proportion of cells along endocycling and endomitosis trajectories following 5 d of treatment with talazoparib (left panels), doxorubicin (center panels), or talazoparib with p21 knockdown (p21KD) (right panels). **t-u**, quantification of nuclear p21 (**t**) and cell state transitions (**u**) after talazoparib or doxorubicin treatment (5 d) in MCF7 (top panels) and T47D cells (bottom panels) administered scrambled (scr) or p21 siRNA. Data in **u** represent mean ± s.d. from three biological replicates. Data in **p-q,t** show representative single-cell distributions for one of three total biological replicates. Talazoparib concentrations, MCF7: 300 nM; T47D: 500 nM. Doxorubicin concentrations, MCF7: 100 nM; T47D: 120 nM.

### Therapy-induced WGD produces a slower growing, drug-resistant polyploid population

Next, we sought to determine the functional consequences of therapy-induced WGD and its impact on the fitness of cancer cells under baseline and drug treatment conditions. After 5 days of talazoparib or doxorubicin treatment, cells were sorted by size to enrich for diploid and polyploid populations and confirmed by measurements of DNA content (**Fig. 5a-b**). Time-lapse imaging and population growth measurements demonstrated that polyploid cells successfully returned to the mitotic cell division cycle to produce proliferative progeny – a cycling polyploid population – after drug removal in both MCF7 and T47D cells (**Fig. 5c, Movie S5**). In all cases, these polyploid cell populations grew more slowly than diploid populations, indicating a proliferative cost for cells that underwent WGD. Similarly, using long-term single-colony imaging, we observed that when drug is removed, diploid cells proliferated rapidly to form large, uniform cell colonies similar to treatment-naive cells, whereas polyploid cells produced progeny at a much slower rate, forming heterogeneous and abnormal colony morphology consisting of cells of varying sizes (**Fig. 5d**). Next, we exposed sorted diploid and polyploid cells to subsequent talazoparib or doxorubicin treatment to evaluate drug sensitivity. We found that polyploid cells were less susceptible to therapy-induced cell cycle arrest than their diploid counterparts or treatment-naive cells (**Fig. 5e-h**). We also observed that polyploid colonies continued to expand at a faster rate than diploid colonies upon retreatment with talazoparib or doxorubicin (**Fig. 5d**). In addition to its cytostatic effects, doxorubicin also elicits potent antitumor activity through the induction of cell death. We observed that doxorubicin induced negative growth rates, indicative of cell death, at lower concentrations in diploid cells in both MCF7 and T47D cells, further demonstrating the drug-resistant properties of the polyploid state (**Fig. 5i-l**). Together, these findings highlight the fitness trade-offs associated with WGD, framing it as a *facultative stress response*: Polyploidization protects against therapy-induced cell cycle arrest and death at the cost of reduced proliferative capacity.

**Fig. 5:**
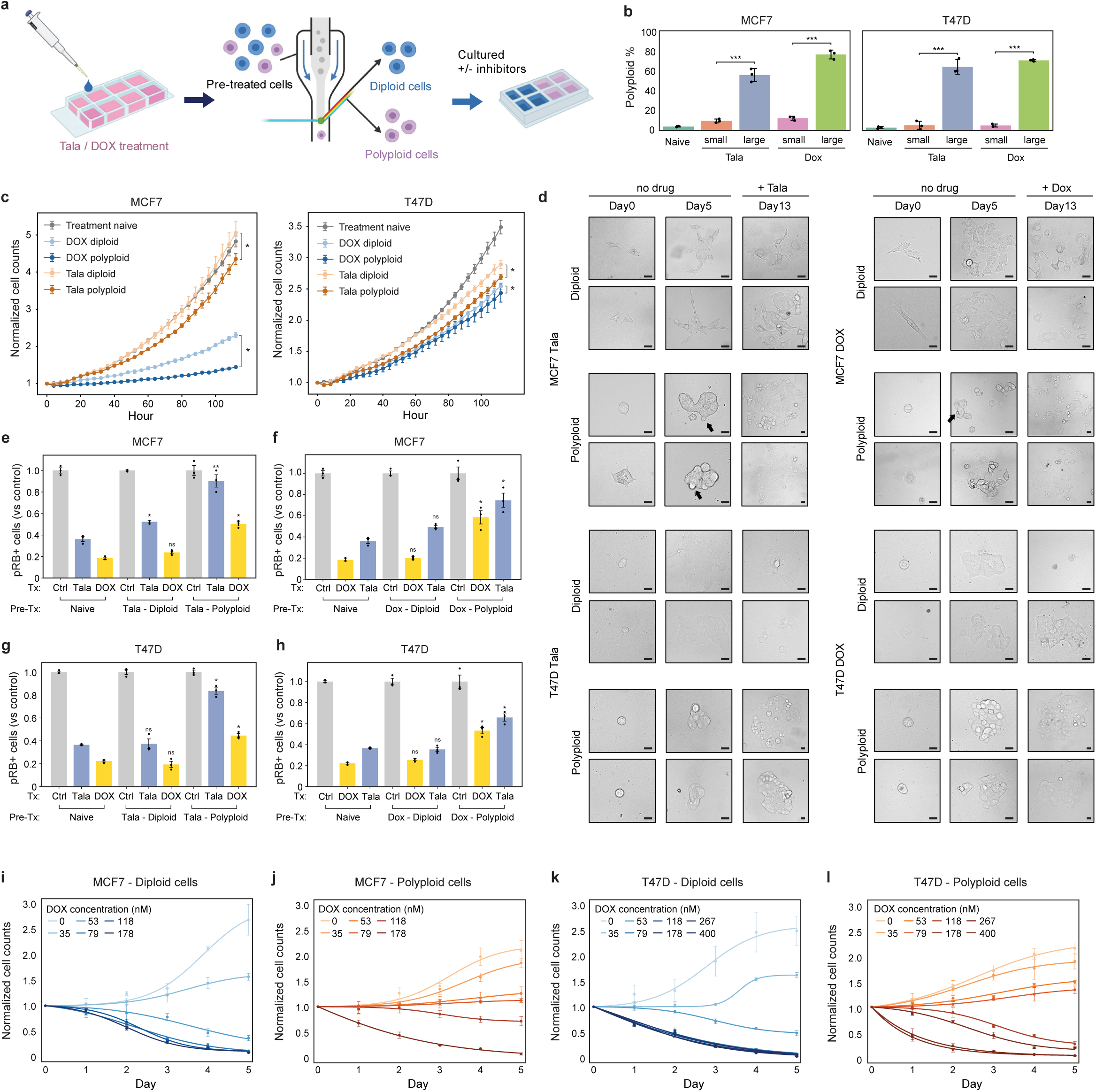
WGD confers resistance to antitumor effects of genotoxic agents. **a-b**, Sorting of polyploid and diploid cells by size (**a**) and quantification of polyploid percentage from IF measurements (**b**) in sorted populations following talazoparib or doxorubicin treatment (5d). **c**, Quantification of cell population normalized to day 0 of treatment-naive and sorted diploid and polyploid cells after drug removal. **d**, Colony growth of sorted diploid and polyploid populations in the absence of drug (days 0 and 5). Talazoparib or doxorubicin was added at day 8 and colonies were reimaged on day 13. **e-h**, Quantification of phospho-RB+ proliferative cells after talazoparib or doxorubicin treatment in drug-naive and sorted diploid and polyploid populations. **i-l**, Quantification of cell population normalized to day 0 in sorted diploid (**i,k**) and polyploid cells (**j,l**) following doxorubicin treatment. In **b-c** and **e-l**, data represent mean ± s.d. from three biological replicates. Statistical significance (*: p < 0.05, **: p < 0.01, ***: p < 0.001) was determined using a two-tailed Student’s *t*-test. Talazoparib concentrations, MCF7: 300 nM; T47D: 500 nM. Doxorubicin concentrations, MCF7: 100 nM; T47D: 120 nM.

### Therapy-induced WGD increases genomic heterogeneity through multiple mechanisms

Prior work demonstrates that WGD can also fuel tumor evolution via genomic instability^62–6667^. In response to talazoparib or doxorubicin treatment, we observed multiple mechanisms driving genomic instability in polyploid cells after WGD. First, cells continue to replicate DNA in the sustained presence of genotoxic agents through endoreplication (**Fig. 1c-e, Extended Data Fig 1d-f**), which itself can produce abnormal karyotypes in the absence of cell division^68^. In addition, polyploid cells possessed elevated γH2AX, phospho-ATR, and 53BP1 compared to diploid cells, a molecular signature indicative of higher levels of replication stress and DNA damage in the endoreplication cycle (**Extended Data Fig. 6**). Upon drug removal, we observed rare occasions of polyploid cells undergoing multipolar mitosis to generate aneuploid progeny (**Extended Data Fig. 7a, Movie S6**). Additionally, both genotoxic agents induced the formation of micronuclei in polyploid cells during endoreplication at a significantly higher frequency than their diploid counterparts (**Fig. 6a-d**). Talazoparib generated nearly twice as many polyploid cells with micronuclei as doxorubicin, highlighting an important functional distinction between PARP1/2 inhibition and topoisomerase II inhibitor chemotherapy. Time-lapse imaging revealed that the mechanism of micronuclei formation depended on treatment context. During talazoparib or doxorubicin treatment, micronuclei were primarily formed by nuclear budding^69–71^ during interphase (**Fig. 6e, Extended Data Fig. 7b, Movie S7**). However, after drug removal, we also observed nuclear fragmentation during cell division as polyploid cells re-entered the mitotic cycle (**Fig. 6f, Extended Data Fig. 7c, Movie S8**). Micronuclei were capable of autonomous cell cycle activity^72^, staining positive for pRB and cyclin A2 even when the primary nucleus or other micronuclei within the same cytoplasm did not (**Extended Data Fig. 7d-e**). Similarly, we found that micronuclei were competent for DNA replication, as indicated by the transient appearance of PCNA foci (**Fig. 6g, Movie S9**). Occasionally, we observed micronuclei being trafficked to the plasma membrane and shed from the mother cell, retaining an intact cytosol and plasma membrane, which generated autonomous aneuploid progeny by amitotic budding, as previously reported^31,32,73,74^ (**Fig. 6h, Extended Data Fig. 7f-g, Movie S9-10**). Altogether, these data demonstrate that the endoreplication cycle is an alternative cancer cell cycle induced by genotoxic agents, capable of producing aneuploid, drug-resistant progeny that can diversify the tumor genomic landscape.

**Fig. 6:**
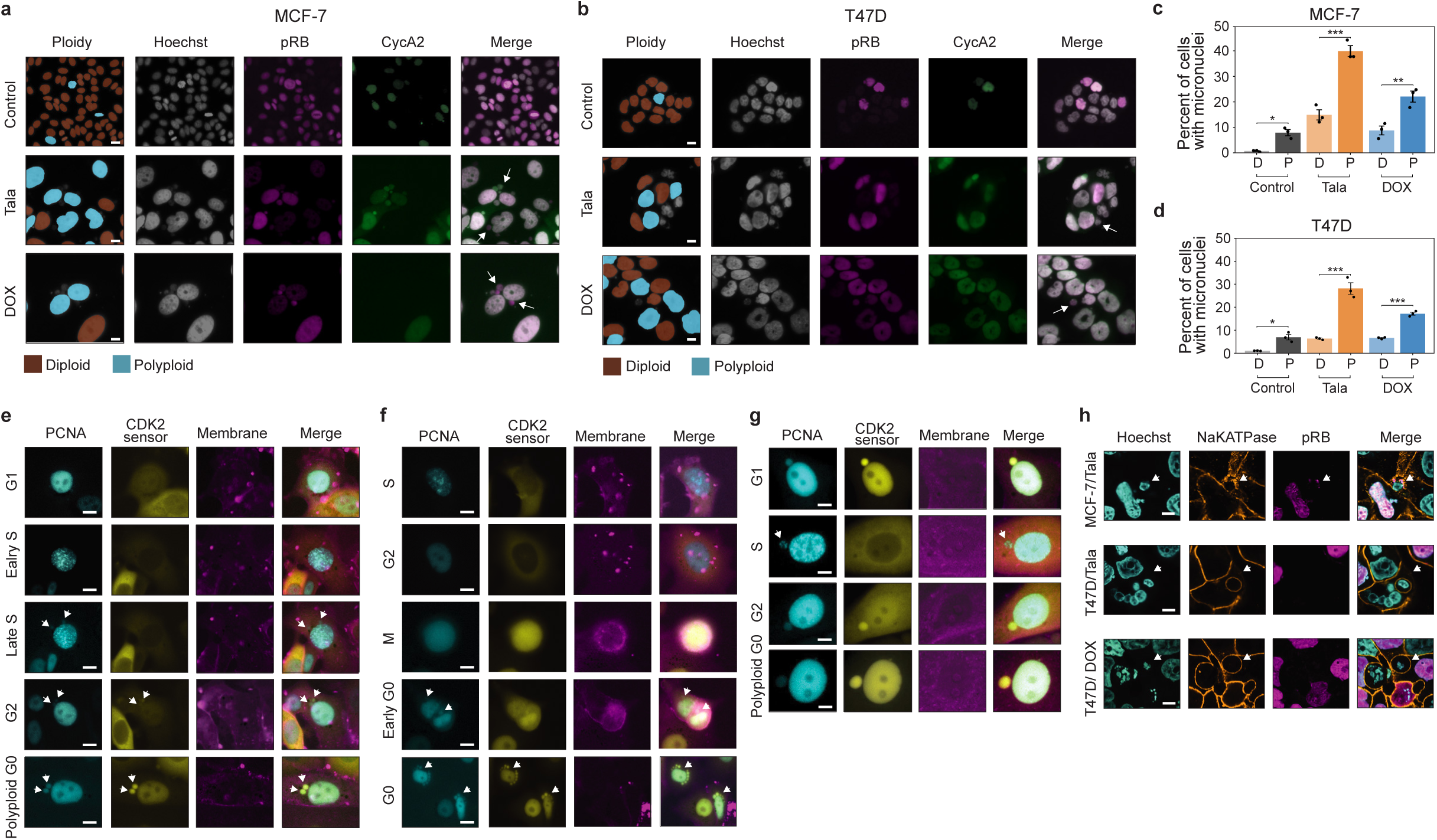
WGD promotes the generation of replication-competent micronuclei and aneuploid microcells. **a-b**, Example IF images of MCF7 (**a**) and T47D cells(**b**) after talazoparib or doxorubicin treatment (5d). Diploid and polyploid labels were assigned based on DNA content, phospho-RB (pRB) and cyclin A2 signatures. White arrows highlight micronuclei that are pRB+ in polyploid cells with primary nuclei that are pRB- and cyclin A2-. **c-d,** Quantification of micronuclei in diploid (D) and polyploid (P) MCF7 (**c**) and T47D cells (**d**). **e-f**, Time-lapse imaging of micronuclei formation in polyploid MCF7 cells (white arrows) following talazoparib treatment during interphase (**e**) and mitosis (**f**). **g**, Time-lapse imaging of micronucleus asynchronously entering S phase in polyploid MCF7 cell (white arrow). **h**, Confocal imaging of microcells formed (white arrows) after talazoparib or doxorubicin treatment in MCF7 and T47D cells. In **c-d**, data represent mean ± s.d. from three biological replicates. Statistical significance (*: p < 0.05, **: p < 0.01, ***: p < 0.001) was determined using a two-tailed Student’s *t*-test. In **a,b,e-g**, scale bars: 20 μM. In **h**, scale bar: 10 μM. Talazoparib concentrations, MCF7: 300 nM; T47D: 500 nM. Doxorubicin concentrations, MCF7: 100 nM; T47D: 120 nM.

### Endocycling and endomitosis converge on a common cyclin D1:CDK4/6-dependent state

Cell cycle mapping suggested that, following cell cycle exit from G2 (endocycling) or M (endomitosis), cells converge onto a single state of cell cycle arrest (polyploid G0) characterized by a loss of RB phosphorylation (**Fig. 4a,f,k**). Subsequent cell cycle re-entry from this G0 state is a key cell state transition in the endoreplication cycle – its regulation serves as a checkpoint that protects against subsequent WGD and its associated genomic instability. To investigate the molecular mechanisms regulating entry from polypoloid G0 to the endoreplication cycle, we trained a random forest classifier to identify cell cycle effectors that distinguish polyploid from diploid G0/G1. Outside of core effectors of the DNA damage response (e.g. γH2AX, 53BP1, p53, p16), cyclin D1 and CDK4 were the most frequently identified features distinguishing cell cycle re-entry in polyploid versus diploid cells across breast cancer cell lines (**Extended Data Fig. 6**). This suggests that activation of the endoreplication cycle may be regulated by cyclin D-mediated activation of CDK4/6 (**Fig. 7a**). Knockdown of cyclin D1 24 h after drug treatment blocked entry into the endoreplication cycle in both MCF7 and T47D cells by trapping cells in polyploid G0 (**Fig. 7b**) and significantly reduced talazoparib- and doxorubicin-induced WGD (**Fig. 7c**). By time-lapse imaging, we also observed that cells transitioning to the endoreplication cycle via endocycling or endomitosis activate CDK4 prior to CDK2 upon cell cycle reentry (**Extended Data Fig. 5h-i**). Consistent with these observations, sequential treatment with the CDK4/6 inhibitor (CDK4/6i) palbociclib 24 hours after DNA damaging agents blocked entry into the endoreplication cycle (**Fig. 7d**) and prevented the acquisition of higher orders of ploidy by WGD in both MCF7 and T47D cells (**Fig. 7e**), further supporting a conserved role for CDK4/6 activation governing the transition to the endoreplication cycle for both endocycling and endomitosis routes to WGD.

**Fig. 7:**
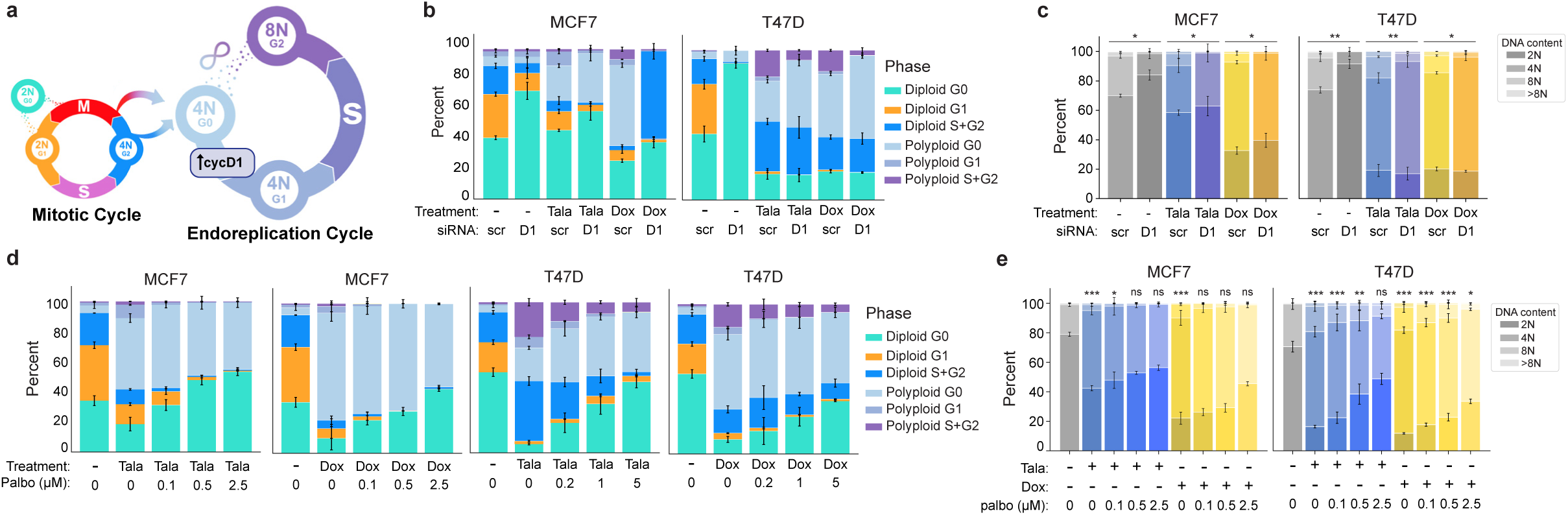
Endocycling and endomitosis converge on a common polyploid G0 state that requires cyclin D1:CDK4/6 activity to enter the endoreplication cycle. a, Schematic of cell state transitions in the mitotic and endoreplication cycle and the mechanisms governing entry to the endoreplication cycle from polyploid G0. **b-c**, Cell state transitions (**b**) and DNA content measurements by single-cell IF (**c**) after talazoparib or doxorubicin treatment (5 d) in MCF7 and T47D cells followed 24 h later by scrambled (scr), or cyclin D1 (D1) siRNA administration. **d-e**, Cell state transitions (**d**) and DNA content measurements by single-cell IF (**e**) after talazoparib or doxorubicin treatment (5 d) in MCF7 and T47D cells followed 24 h later by palbociclib (palbo) at the indicated concentrations. In **b-e**, data represent mean ± s.d. from three biological replicates. Statistical significance (*: p < 0.05, **: p < 0.01, ***: p < 0.001) was determined using a two-tailed Student’s *t*-test (**c**) and a one-way ANOVA with Tukey’s *post hoc* test (**e**). Talazoparib concentrations, MCF7: 300 nM; T47D: 500 nM. Doxorubicin concentrations, MCF7: 100 nM; T47D: 120 nM.

### Targeting the endoreplication cycle sensitizes cells to genotoxic agents and reduces therapy-induced genomic instability

DNA-damaging therapies trigger cell death and induce exit from the mitotic cell division cycle in actively proliferating cells. We have shown that exit from the mitotic cell cycle is not a permanent state of cell cycle arrest as cells can switch to a drug-tolerant endoreplication cycle in a cyclin D1:CDK4/6-dependent manner. Based on these findings, we postulated that blocking entry to the endoreplication cycle by targeting CDK4/6 activation should permit the antitumor effects of genotoxic therapies while preventing subsequent WGD to preserve genome integrity. However, concurrent treatment of talazoparib or doxorubicin with palbociclib protected cells from therapy-induced cell death (**Fig. 8a-b**), likely due to the cytostatic effects of palbociclib itself. On the other hand, sequential treatment with doxorubicin for 24h before palbociclib treatment successfully preserved the cytotoxic effect of doxorubicin (**Fig. 8a-b**). Similarly, the growth of patient-derived organoids (PDOs) derived from triple-negative breast cancer (TNBC) was significantly more attenuated in sequential versus combination therapy, inducing a similar growth retardation as talazoparib or doxorubicin treatment alone (**Fig. 8c-d**). In colony-formation assays, sequential palbociclib treatment also significantly reduced the outgrowth of clones compared to concurrent therapy with talazoparib or doxorubicin in MCF7 and T47D cells (**Fig. 8e-f**). Although 10 d of sustained treatment with genotoxic agents alone prevented the outgrowth of individual colonies, the clones that survived reentered the mitotic cycle upon drug removal to generate proliferative (pRB+) colonies with very high levels of ploidy (**Fig. 8h-i**). This indicates that the drug-tolerant persistor cells successfully transitioned to the endoreplication cycle as a mechanism to survive prolonged genotoxic stress before switching back to the mitotic cycle to generate a proliferative polyploid population upon drug removal. We also detected extensive γH2AX staining and cytoplasmic DNA in these proliferative polyploid colonies (**Fig. 8h-i**). This is consistent with micronucleus rupture^75^ and indicates significant genomic instability in surviving colonies. Moreover, we found that sequential therapy with palbociclib also inhibited the initial generation of micronuclei in both MCF7 and T47D cells (**Fig. 8j-l**), further positioning CDK4/6 inhibitors as effective WGD-blocking agents that can decouple the cytotoxic efficacy of DNA-damaging agents from the undesirable genomic instability that typically accompanies their use.

**Fig. 8:**
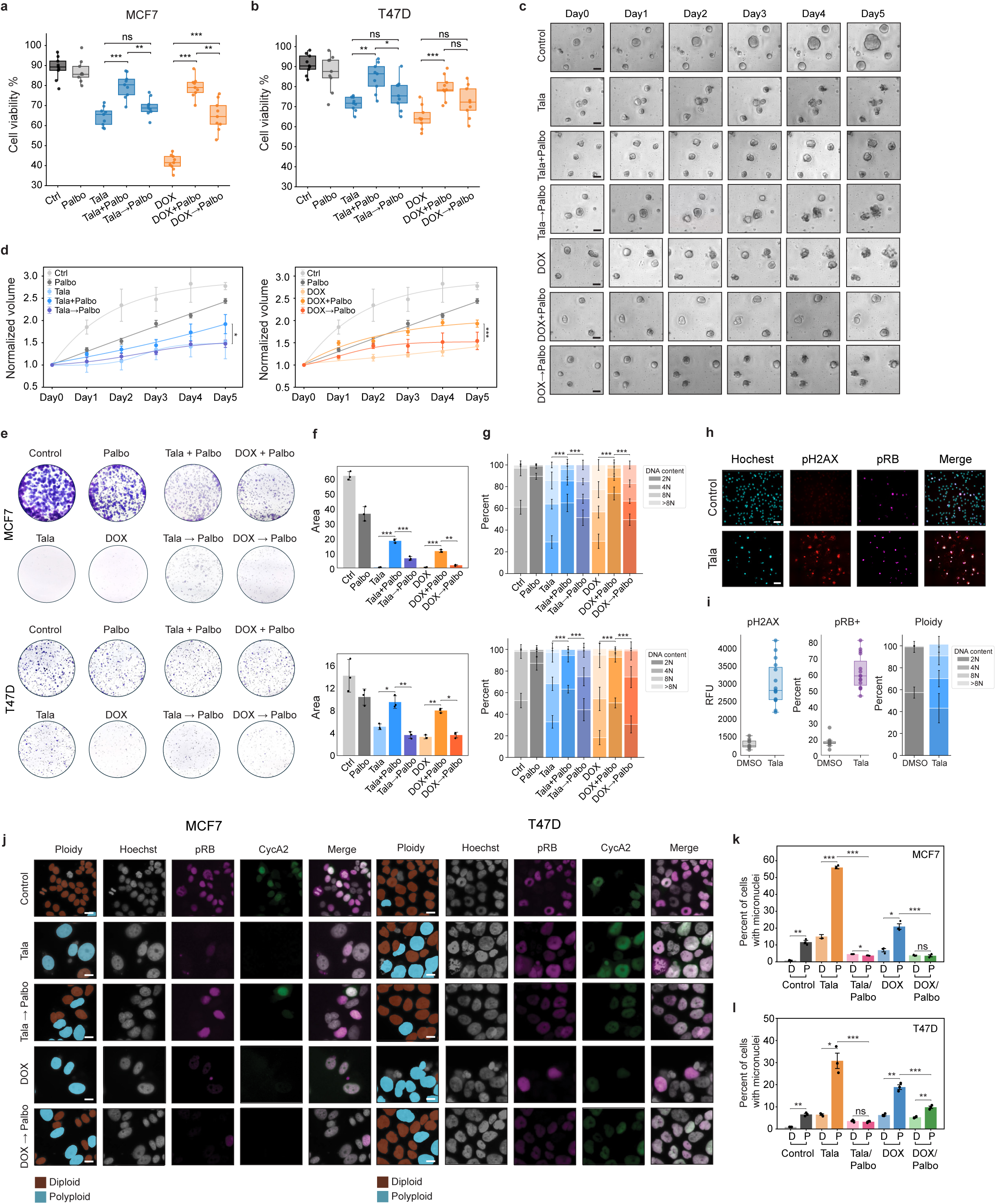
Sequential treatment with DNA-damaging agents and CDK4/6 inhibitors preserves antitumor response while blocking genomic instability. **a-b**, Cell viability by trypan blue staining in MCF7 (**a**) and T47d cells (**b**). **c-d**, Example images (**c**) and quantification of growth of patient-derived organoids (**d**) following 5 d of concurrent or sequential treatment with talazoparib/doxorubicin and palbociclib. Growth is calculated as the fold-change increase in organoid volume compared to day 0. **e-g**, Example images (**e**), total colony area (**f**), and colony ploidy (**g**) following 10 d of concurrent or sequential treatment with talazoparib/doxorubicin and palbociclib. **h-i**, Example images (**h**), and quantification of phospho-H2AX, phospho-RB, and ploidy (**i**) by single-cell IF in colonies treated for 10 d with talazoparib or doxorubicin followed by 20 d of drug removal. **j-l**, Example images of MCF7 and T47D cells (**j**) and quantification of micronuclei formation (**k-l**) after concurrent or sequential treatment with talazoparib/doxorubicin and palbociclib. In **a-b, f-g, i, k-l**, data represent mean ± s.d. from three biological replicates. In **d**, data represent population means ± s.e.m. from three biological replicates (n=62-259 individual organoids per condition). Statistical significance (*: p < 0.05, **: p < 0.01, ***: p < 0.001) was determined using a one-way ANOVA with Tukey’s *post hoc* test (**a-b, f-g, k**) or a two-tailed Student’s *t*-test (**d**). In **c,j**, scale bars: 20 μM. In **h**, scale bars: 80 μM. Talazoparib concentrations, MCF7: 300 nM; T47D: 500 nM. Doxorubicin concentrations, MCF7: 100 nM; T47D: 120 nM. Palbociclib concentrations, MCF7: 374 nM; T47D: 1 μM.

## DISCUSSION

Whole-genome duplication (WGD) represents a pivotal event in the life history of a cancer cell, acting as both a buffer against lethal genetic imbalances and a catalyst for rapid genomic evolution. In this paper, we show that therapy-induced polyploidization allows cancer cell populations to persist under stressful conditions and provides a reservoir of genomic diversity to accelerate adaptation. We demonstrate that breast cancer cells exposed to distinct classes of genotoxic agents (PARP1/2 and topoisomerase II inhibitors) promote the transition from the mitotic cell division cycle to the endoreplication cycle through two distinct trajectories governed by p21 dynamics: (1) cell cycle exit in G2 (endocycling) and (2) mitotic entry and subsequent abortion (endomitosis). Regardless of the route taken, these paths converge on a mononuclear, CDK4/6-dependent state of arrest. This convergence suggests a shared molecular vulnerability to therapy-induced WGD that can be targeted through the precise temporal sequencing of genotoxic and WGD-blocking agents. Our results also highlight a crucial fitness trade-off associated with therapy-induced WGD^33,36,76,77^. On one hand, the polyploid state confers resistance to genotoxic stress (**Fig. 5e-l**) and provides fuel for tumor evolution through augmented genomic instability, including increased replication stress (**Fig. 8h-i**, **Extended Data Fig. 6a**), extensive micronuclei formation (**Fig. 6a-f, Extended Data Fig. 7b-e**), and the amitotic budding of aneuploid microcells (**Fig. 6g-h, Extended Data Fig. 7f-g**). On the other hand, these adaptations come at a fitness cost of reduced proliferative capacity (**Fig. 5c-d**). This frames therapy-induced WGD as a facultative stress response in which cancer cells transiently shift from the canonical mitotic cell cycle to endoreplication to withstand and adapt to stressful environments, and subsequently revert back to the mitotic cell cycle once stress subsides.

The distinction between endocycling and endomitosis has traditionally been studied in the context of developmental biology^78,79^. Our observation that these disparate cell cycle variants can co-exist within the same clonal population in response to identical stress suggests a high degree of plasticity in the DNA damage response (DDR). We identify p21 as the molecular switch governing this bifurcation. The induction of p21 during G2 promotes endocycling by effectively blocking CDK activity and triggering cell cycle exit. An insufficient p21 response results in an extended but “leaky” G2 arrest, from which cells can slip into a prolonged and ultimately unsuccessful mitosis, leading to the reformation of a single nucleus and cell cycle exit from M phase via endomitosis. This model helps reconcile conflicting reports regarding the necessity of p53 for WGD^27^. Although p53 is the primary driver of p21 expression in the DDR, our data from p53-mutant cell lines (e.g., T47D) indicate that residual p53 activity or non-canonical mechanisms of p21 induction^80–82^ can still drive endocycling, albeit with lower efficiency. Thus, the route to WGD is not determined solely by the binary status of p53, but by the quantitative capacity of the cell to reach a p21 threshold during the G2 DDR.

We show that both routes of mitotic cell cycle exit converge on a common polyploid G0 state that requires the activation of cyclin D1:CDK4/6 to complete the transition to the endoreplication cycle. Historically, CDK4/6 inhibitors such as palbociclib are viewed as agents that arrest diploid cells in a G0 state. Our results expand this paradigm by identifying CDK4/6 as the gatekeeper of the endoreplication cycle, consistent with recent reports that cells transit through a CDK4/6-dependent state during ribotoxic stress-induced WGD^82^. This dependence creates a specific vulnerability: Although polyploid cells exhibit enhanced resistance to the DNA-damaging agents that produced them, they remain sensitive to CDK4/6 inhibitors, which function as WGD-blocking agents that attenuate therapy-induced genomic instability and the acquisition of stable solutions to drug resistance. These biological insights have translational implications. We show that concurrent administration of CDK4/6i with PARP1/2 or topoisomerase II inhibitors is counterproductive: By arresting cells in diploid G0, CDK4/6i prevents the replication stress during S phase that is required for the cytotoxic efficacy of these genotoxic drugs. Conversely, sequential therapy – treating with DNA-damaging agents followed by CDK4/6 inhibitors — leverages cell cycle biology to maximize cellular response. It allows the genotoxic therapy to kill a bulk population of tumor cells while subsequently locking the surviving cells in a polyploid G0 state, preventing the therapy-induced genomic instability associated with WGD. While we demonstrate the efficacy of this therapeutic strategy in PDOs from breast tumors (**Fig. 8c-d**) and colony-forming assays (**Fig. 8e-g**), additional *in vivo* studies comparing pulsed or sequential dosing schedules versus continuous combination strategies are required to validate these findings further in a preclinical setting. However, our findings are consistent with and provide a mechanistic rationale to explain the positive clinical benefit observed when CDK4/6 inhibitors are administered as maintenance therapy following chemotherapy in breast and ovarian cancers^83–85^.

In summary, we elucidate the molecular mechanisms governing the therapy-induced switch from the mitotic to the endoreplication cycle in breast cancer. By revealing the paths cells take through cell cycle state space in response to DNA-damaging agents and the molecular mechanisms that govern them, we provide a roadmap for the rational design of combination therapies that not only trigger tumor cell arrest or death, but also block the evolutionary escape routes that lead to drug resistance and recurrence.

## EXTENDED DATA FIGURE LEGENDS

**Extended Data Fig. 1:**
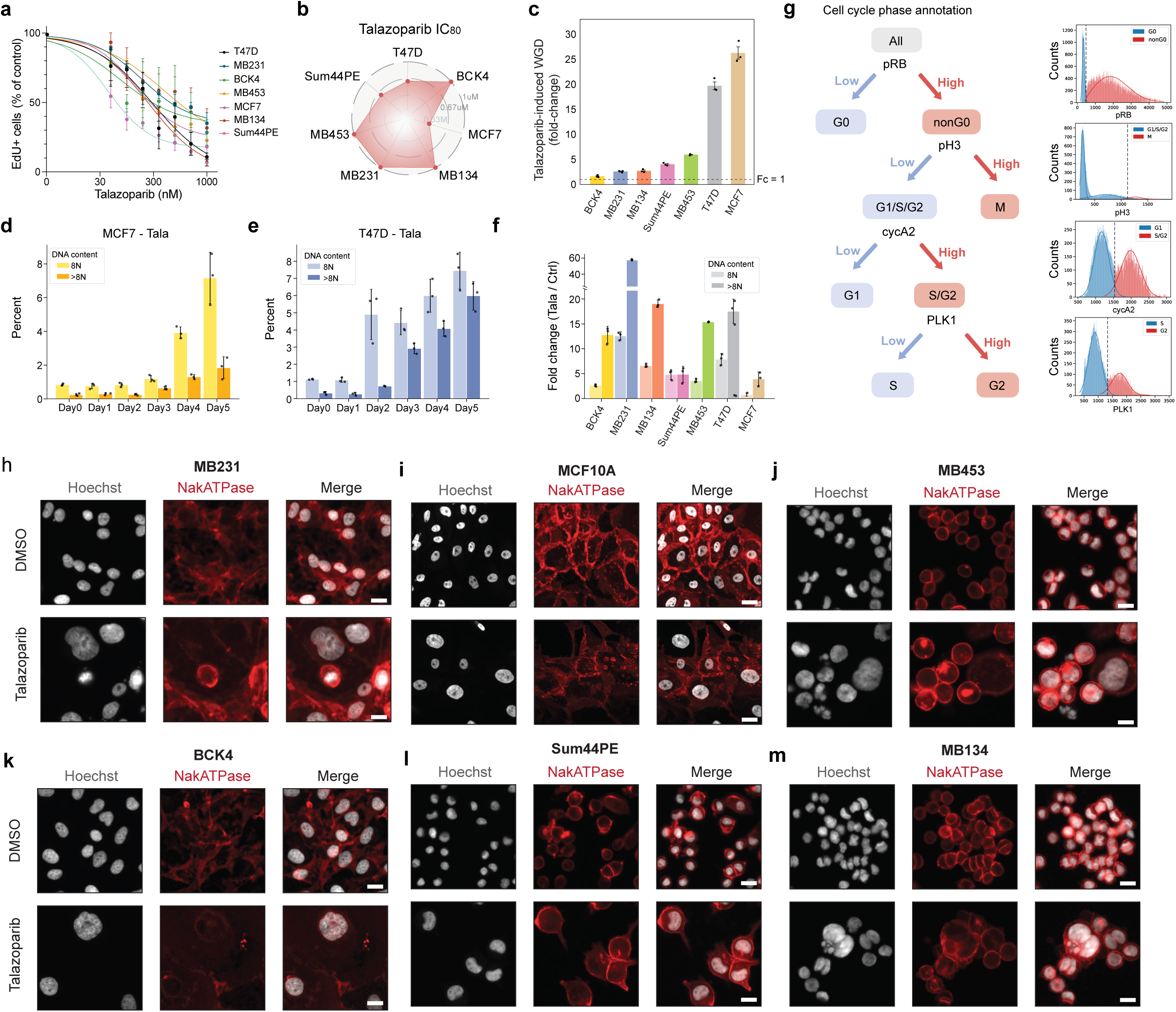
**a-b**, EdU positivity by single-cell IF in response to increasing concentrations of talazoparib (5 d) in a panel of breast cancer cell lines (**a**) and potency (IC_80_) calculations (**b**). **c**, Fold-change increases in talazoparib-induced WGD. **d-f**, Time-course of the generation of polyploid cells with 8N and >8N DNA content after talazoparib treatment in MCF7 (**d**), T47d (**e**), and after 5 d of talazoparib treatment in cell line panel (**f**). **g**, Schematic of cell cycle phase annotation strategy. **h-m**, Example IF images of control and talazoparib-treated cell lines (5 d); scale bar: 20 μM. In **a,c,d-f**, data represent mean ± s.d. from three biological replicates. In **c-f, h-m,** talazoparib was administered at IC_80_ concentrations for each cell line.

**Extended Data Fig. 2:**
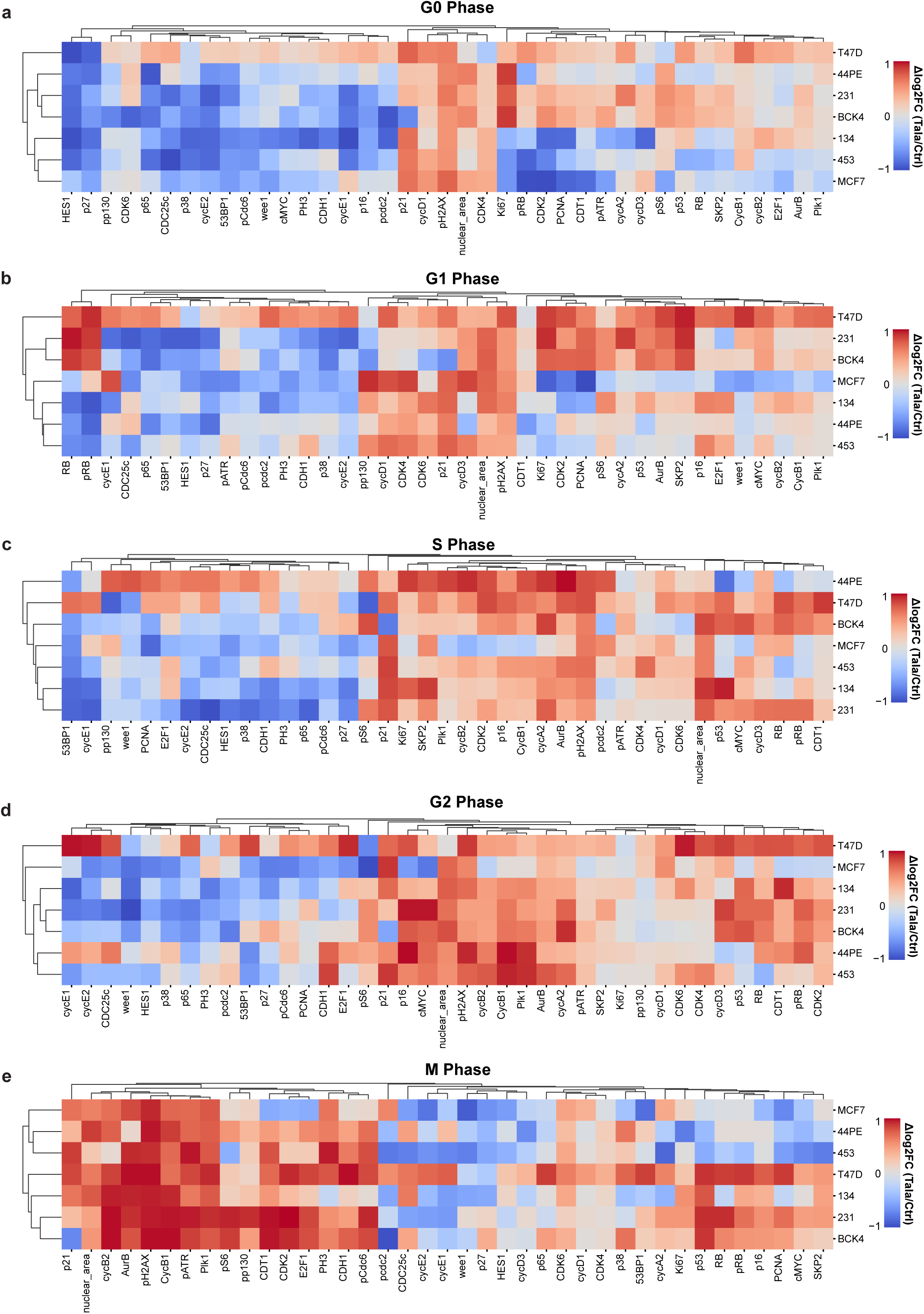
**a-b**, Heatmaps of talazoparib-induced proteomic profiles across cell line panel per cell cycle phase derived from multiplexed IF data.

**Extended Data Fig. 3:**
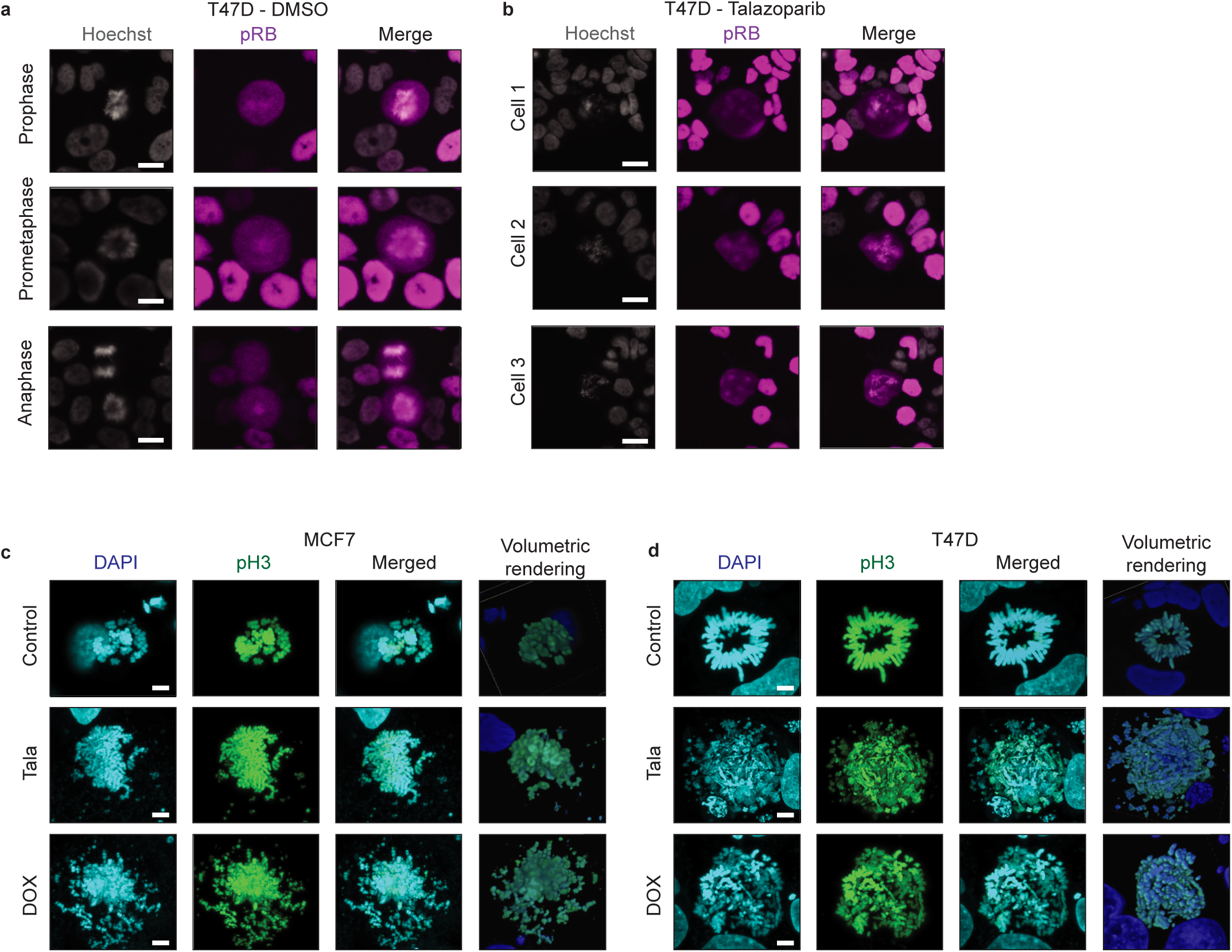
**a-b**, Example IF images of control (**a**) and talazoparib-treated (**b**) T47D cells showing normal (**a**) and perturbed spindle organization (**b**), including multipolar spindle formation (Cells 1-3) and centrosome clustering (Cell 1). **c-d**, Example maximum intensity projections (DAPI/phospho-H3/Merged) and volumetric rendering of confocal IF images of control and talazoparib-treated MCF7 (**a**) and T47D cells (**b**). In **a-b**, scale bars: 20 μM. In **c-d**, scale bars: 5 μM.

**Extended Data Fig. 4:**
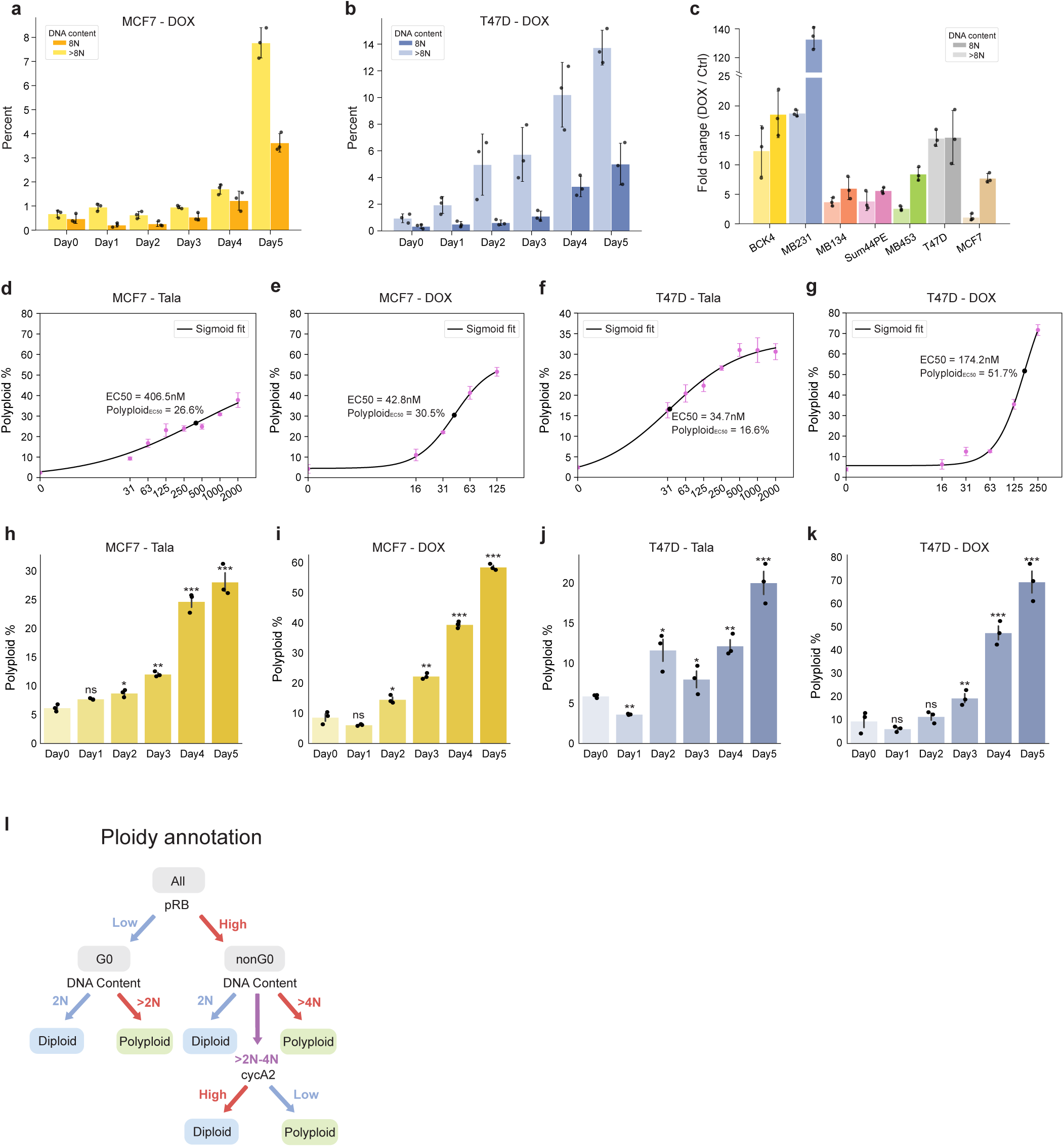
**a-c**, Time-course of doxorubicin-induced polyploidy in MCF7 (**a**), T47D (**b**), and after 5 d of treatment in other cell lines (**c**). **d-g**, Concentration-response curves of talazoparib- and doxorubicin-induced polyploidy in MC7 (**d-e**) and T47D cells (**f-g**). **h-k**, Time-course of talazoparib- and doxorubicin-induced polyploidy in MCF7 (**h-i**) and T47D cells (**j-k**). **i**, Schematic of ploidy annotation strategy. In **a-k**, data represent mean ± s.d. from three biological replicates. In **h-k**, statistical significance (*: p < 0.05, **: p < 0.01, ***: p < 0.001) was determined using a one-way ANOVA with Tukey’s *post hoc* test comparing Days 1-5 to Day 0.

**Extended Data Fig. 5:**
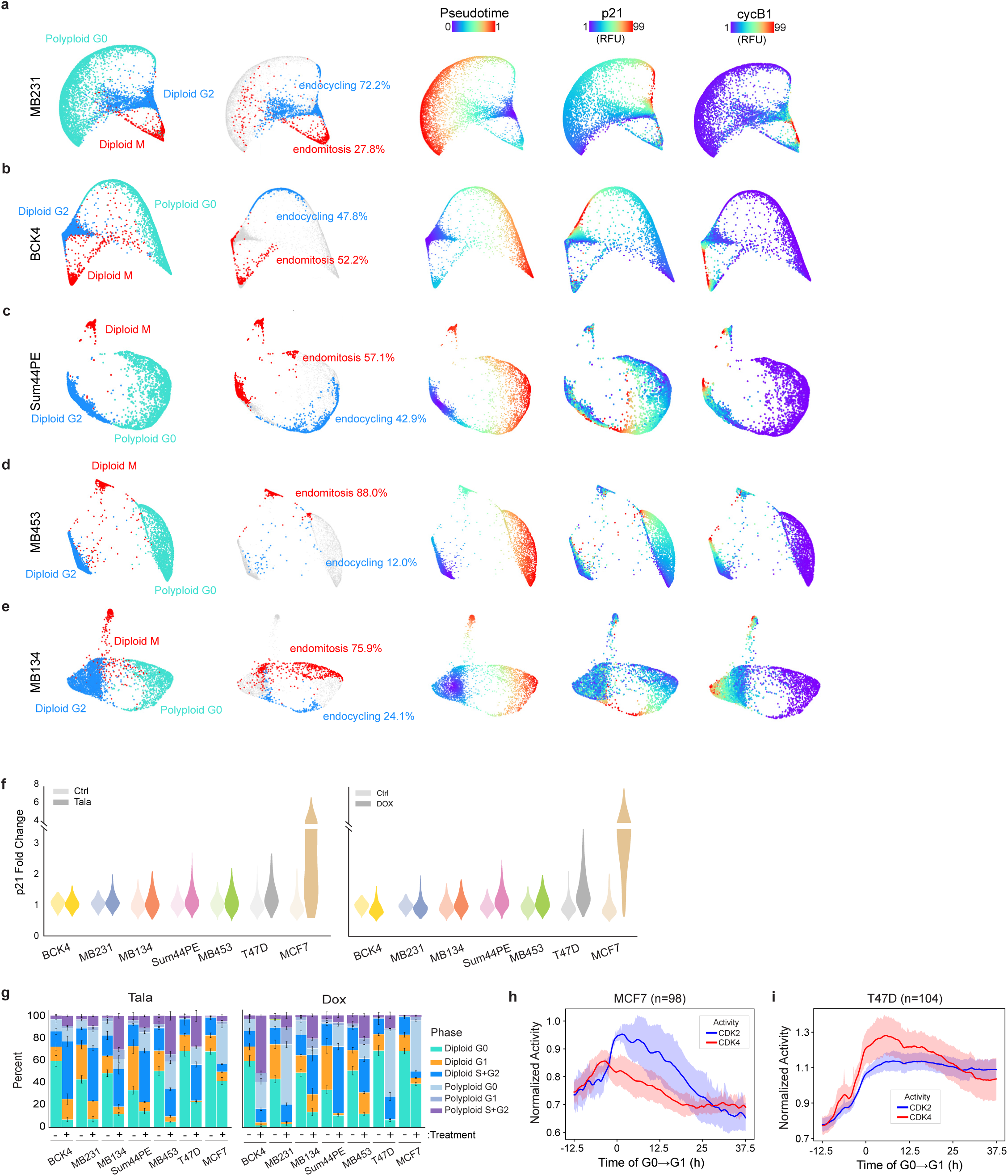
**a-e**, Cell cycle maps of MDA-MB-231 (**a**), BCK4 (**b**), Sum44PE (**c**), MDA-MB-453 (**d**), MDA-MB-134 cells (**e**) showing bifurcation of diploid G2 cell into diploid M and polyploid G0, Leiden-based clustering and quantification of cells along the endocycling and endomitosis trajectories, pseudotime values, nuclear p21, and nuclear cyclin B1 intensities. **f**, Quantification of p21 induction (fold-change versus control) after talazoparib (left panel) or doxorubicin (right panel) treatment in cell lines. **g**, Cell state transitions in cell line panel in response to 5 d treatment with IC_80_ concentrations of talazoparib and doxorubicin. **h-i**, Average CDK2 (blue) and CDK4 activity of polyploid G0 MCF7 (**h**) and T47D cells (**i**) synchronized *in silico* at time of G0 to G1 transition. Lines: average of n=98 (**h**) and n=104 (**i**) individual cell traces. Shaded areas: 95% CI.

**Extended Data Fig. 6:**
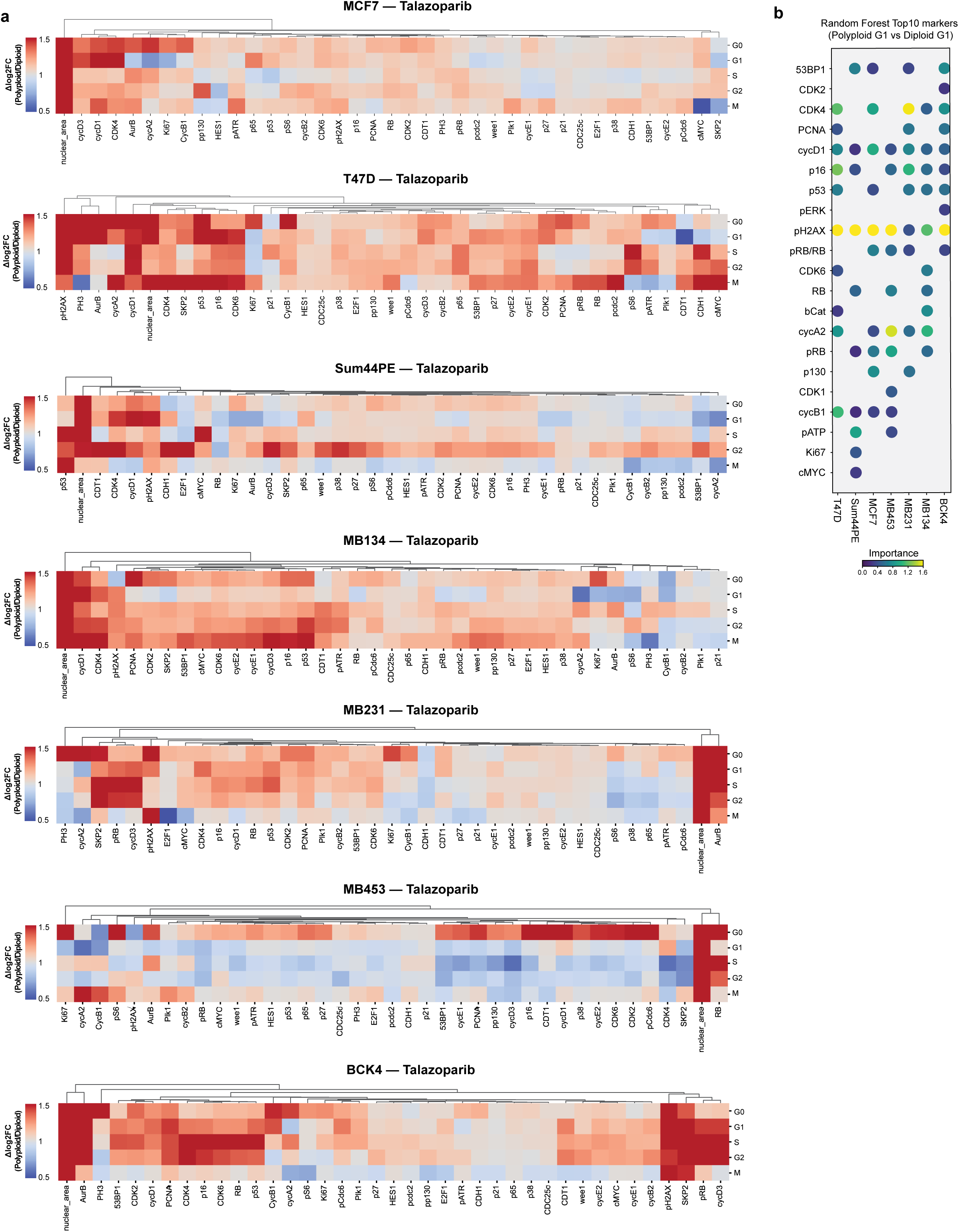
**a**, Fold-change enrichment of proteins by cell cycle phase in polyploid versus diploid cells after 5 d treatment with IC_80_ concentrations of talazoparib derived from multiplexed IF data. **b**, Variable importance calculations of top 10 proteins distinguishing polyploid from diploid G1 cells following talazoparib treatment in cell line panel. Dot color represents variable importance. Only features that appeared in the top 10 of at least one cell line are shown.

**Extended Data Fig. 7:**
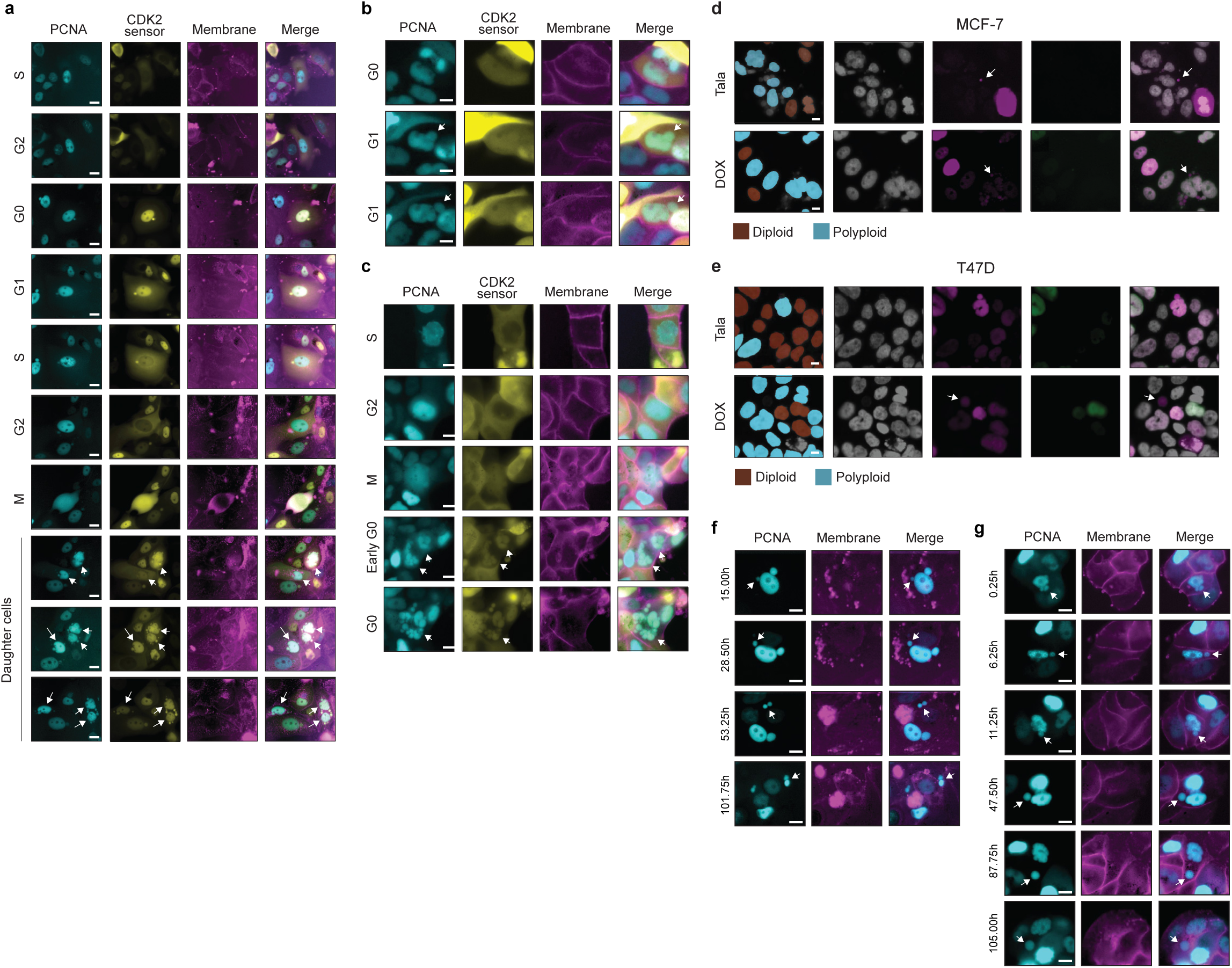
**a**, Time-lapse imaging of multipolar mitosis in polyploid MCF7 cells generating aneuploid progeny (white arrows) after talazoparib removal (**a**). **b-e**, Time-lapse imaging of micronuclei formation (white arrows) in polyploid T47D cells following talazoparib treatment during interphase (**b**) and mitosis (**c**) and examples of pRB+ micronuclei in MCF7 (**d**) and T47D cells (**e**) where the primary nucleus is pRB-. **f-g**, Time-lapse imaging of microcell budding (white arrows) in MCF7 (**d**) and T47D cells (**e**) during talazoparib treatment. Scale bars: 20 μM.

## SUPPLEMENTIAL MOVIE LEGENDS

**Movie S1:** Time-lapse imaging of talazoparib-induced endocycling in MCF7 cells. Top panels display PCNA-mTurqoise2, DHB-mVenus (CDK2 activity sensor), and CDK4-KTR-mCherry. Bottom panels show the corresponding single-cell time courses of nuclear PCNA intensity, CDK2 activity, and CDK4 activity.

**Movie S2:** Time-lapse imaging of T47D cells treated with talazoparib, illustrating prolonged G2 arrest. Top panels display PCNA-mTurqoise2, DHB-mVenus (CDK2 activity sensor), and CDK4-KTR-mCherry. Bottom panels show the corresponding single-cell time courses of nuclear PCNA intensity, CDK2 activity, and CDK4 activity.

**Movie S3:** Time-lapse imaging of talazoparib-induced endocycling in T47D cells. Top panels display PCNA-mTurqoise2, DHB-mVenus (CDK2 activity sensor), and CDK4-KTR-mCherry. Bottom panels show the corresponding single-cell time courses of nuclear PCNA intensity, CDK2 activity, and CDK4 activity.

**Movie S4:** Time-lapse imaging of talazoparib-induced endomitosis in T47D cells. Top panels display PCNA-mTurqoise2, DHB-mVenus (CDK2 activity sensor), and CDK4-KTR-mCherry. Bottom panels show the corresponding single-cell time courses of nuclear PCNA intensity, CDK2 activity, and CDK4 activity.

**Movie S5:** Time-lapse imaging of MCF7 cells during 5 d of talazoparib treatment, showing generation of a polyploid cell by endocycling followed by mitotic cell division of polyploid cell after drug removal. Left panel: PCNA-mTurqoise2, Right panel: DHB-mVenus (CDK2 activity sensor). The timing of drug treatment and removal is indicated in the upper left corner and cell cycle phase transitions are annotated.

**Movie S6:** Time-lapse imaging of MCF7 cells during 5 d of talazoparib treatment, showing generation of a polyploid cell by endocycling followed by multipolar mitosis after drug removal. Left panel: PCNA-mTurqoise2, Right panel: DHB-mVenus (CDK2 activity sensor). The duration of drug treatment is indicated in the upper left corner and cell cycle phase transitions are annotated.

**Movie S7:** Time-lapse imaging of MCF7 cells during 5 d of talazoparib treatment, showing generation of a polyploid cell by endocycling followed by micronuclei formation by nuclear budding during interphase. Left panel: PCNA-mTurqoise2, Right panel: DHB-mVenus (CDK2 activity sensor). The duration of drug treatment is indicated in the upper left corner and cell cycle phase transitions are annotated.

**Movie S8:** Time-lapse imaging of MCF7 cells during 5 d of talazoparib treatment, showing generation of a polyploid cell by endocycling followed by micronuclei formation during mitosis. Left panel: PCNA-mTurqoise2, Right panel: DHB-mVenus (CDK2 activity sensor). The duration of drug treatment is indicated in the upper left corner and cell cycle phase transitions are annotated.

**Movie S9:** Time-lapse imaging of polyploid MCF7 cells after 5 d of talazoparib treatment, showing micronucleus that enters S phase, as indicated by transient PCNA foci, followed by amitotic budding and release of autonomous microcell. Merged images of PCNA-mTurqoise2 (cyan) and CellBright Steady plasma membrane dye (magenta).

**Movie S10:** Volumetric rendering of z-stack of polyploid T47D cell and associated microcell. Merged images of Hoechst (cyan) and NaKATPase (orange, as a plasma membrane marker).

## METHODS

### Cell culture

Seven breast cancer cell lines (MDA-MB-231, MDA-MB-453, MC7, T47D, MDA-MB-134-VI, SUM-44PE, and BCK4) were cultured at 37 °C in a humidified atmosphere containing 5 % CO₂. The detailed base media, serum sources, and supplement compositions for each cell line are summarized in **Table S1**.

### Lentivirus production and transduction

Stable cell lines were established by lentiviral transduction of pLenti-DHB-mVenus-p2a-mCherry-CDK4KTR (Addgene, Plasmid #126679) and pLenti-PGK-Puro-TK-NLS-mTurq2-PCNA (Addgene, Plasmid #118617). For lentivirus production, HEK293FT cells were co-transfected with each biosensor together with psPAX2 (Addgene, Plasmid #12260) and pMD2.G (Addgene, Plasmid #12259) using Lipofectamine 2000 (Invitrogen) in OptiMEM. Viral supernatants were harvested 72 h post-transfection, filtered through a 0.45 µm filter, and stored at –80 °C until use. For transduction, target cells were incubated with viral supernatants in complete medium containing 6 µg/mL polybrene. Cells were incubated 24h before replacing the medium with fresh culture medium. Drug selection was initiated 48–72 h post-infection for constructs containing puromycin resistance. Following antibiotic selection, fluorescence-activated cell sorting (FACS) was performed to isolate cell populations co-expressing biosensor fluorescence signals.

### EdU incorporation assay

DNA synthesis was assessed using the Click-iT™ EdU Alexa Fluor™ 488 Imaging Kit (Thermo Fisher Scientific) according to the manufacturer’s protocol with minor modifications. Cells were seeded onto 96-well glass-bottom plates (Cellvis) pre-coated with bovine plasma fibronectin (Sigma-Aldrich) at 1 µg/cm² for 30 min at 37 °C to enhance cell attachment. Cells were treated with the indicated drugs - talazoparib (MedChemExpress, MCE), doxorubicin (MCE), or palbociclib (MCE) - at the concentrations specified. Drug treatments were maintained for five days, after which 10 µM EdU was added (4h, 37 °C, 5% CO₂). After labeling, cells were fixed with 4 % paraformaldehyde for 30 min at room temperature and permeabilized with 0.3 % Triton X-100 (Sigma-Aldrich) for 30 min. The Click-iT reaction cocktail was applied for 30 min in the dark, and nuclei were counterstained with Hoechst 33342 (5 µg/mL, 15 min). Plates were mounted in 50% glycerol/PBS before imaging. Fluorescence images were acquired using the Leica Thunder inverted fluorescence microscope using the HC PL APO 20x/0.80 objective and K8 sCMOS camera. The percentage of EdU-positive cells was quantified based on Cellpose 3.0^86^ segmentation followed by regionprops-based quantification, as described in the Data analysis section.

### siRNA transfection

Transient knockdown experiments were performed using Lipofectamine RNAiMAX transfection reagent (Thermo Fisher Scientific, Cat. No. 13778075) according to the manufacturer’s protocol. Cells were seeded into 96-well plates to achieve 40–60% confluence at the time of transfection. For each well, siRNA (cyclin D1: Dharmacon, L-003210-00-0005, final concentration 8 nM; p21: Dharmacon, L-003471-00-0005, final concentration 25 nM) was diluted in Opti-MEM (Thermo Fisher Scientific, Cat. No. 31985070), mixed with Lipofectamine RNAiMAX, and incubated for 20 min at room temperature. After 48-96h of incubation, cells were fixed, and knockdown was quantitatively assessed by immunofluorescence imaging. A non-targeting scrambled siRNA pool (Dharmacon, Cat. No. D-001810-10-05) was used as a negative control.

### Fluorescence-activated cell sorting (FACS)

Cells were expanded to approximately 2–5 x 10^7^ prior to sorting. Following dissociation, cells were resuspended in PBS supplemented with 2% FBS and 1% penicillin–streptomycin, filtered through a 0.45 μm strainer to remove aggregates, and maintained on ice until analysis. Sorting was performed using the Bigfoot Spectral Cell Sorter (ThermoFisher Scientific) under sterile conditions, and sorted cells were collected into tubes containing fresh medium. For the separation of diploid versus polyploid cells, forward scatter was used as a proxy for cell size, and discrete gates were applied to isolate smaller (diploid-enriched) and larger (polyploid-enriched) fractions.

### *In situ* paraffin embedding

To aid cell adhesion during the multiplexed immunofluorescence procedure (see below), formalin-fixed paraffin-embedded (FFPE) chamber slides were prepared following an optimized protocol. Cell lines grown on removable chambered slides (ibidi, Cat. No. 80841) were rinsed with PBS (300 µL/well) and fixed in 4% paraformaldehyde for 24 h at room temperature. The chambers were removed before dehydration. Slides were sequentially immersed in 30%, 50%, 70%, 95%, and 100% ethanol (each for 1h, with 70% ethanol optionally overnight at 4 °C), followed by two xylene incubations (1h each) and overnight paraffin infiltration at 60°C. Slides were then cooled at room temperature for at least 24 h before further processing. Slides were positioned vertically on a rack and dewaxed at 60°C until the paraffin was completely melted plus an additional 2h to ensure complete removal of residual paraffin. For deparaffinization and rehydration, slides were immersed twice in xylene (5 min each), followed by graded ethanol washes (100%, 90%, 80%, and 70%, 5 min each) and rinsed in PBS.

### Multiplexed Immunofluorescence Imaging

Chamber slides prepared as described above were permeabilized with 0.3% Triton X-100 in PBS for 30 min at room temperature. Antigen retrieval was performed following the Leica CellDive manufacturer’s instructions, using the recommended buffers and heating conditions. After retrieval, slides were washed thoroughly in PBS and mounted into chamber wells for downstream staining. For immunostaining, slides were incubated in blocking buffer (3% BSA/10% donkey serum in PBS) for 1 h at room temperature, followed by incubation with primary antibodies diluted in 3% BSA/PBS for 1 h on a rocker. After three washes with PBS, slides were incubated with Hoechst for 15 min, rinsed again in PBS, and mounted in 50% glycerol in PBS. Multiplexed imaging was performed on the Leica CellDive system, using iterative cycles of antibody staining, imaging, and fluorophore inactivation. After each imaging round, fluorophores were inactivated by incubating slides in 10% H₂O₂ + 20% NaHCO₃ freshly prepared in ddH₂O for 15 min at room temperature, followed by a 5 min PBS wash and Hoechst recharging for 5 min. All antibodies and fluorophores used in this study are listed in Table S2. Subsequent image processing, quantification, and generation of cell cycle maps are described in the Data analysis section.

### Live-cell membrane labeling

For live-cell membrane labeling during time-lapse imaging, cells were stained using the CellBrite Steady Membrane Staining Kit (Biotium). Briefly, cells were incubated with 1× Dye at 37 °C for 30 min, followed by incubation with 1× Enhancer at 37 °C for an additional 30 min. After staining, cells were washed and maintained in medium supplemented with 0.2× Dye and 0.1× Enhancer for live-cell imaging. During prolonged time-lapse experiments, imaging medium containing 0.2× Dye and 0.1× Enhancer was refreshed every three days to maintain membrane signal intensity.

### Time-lapse imaging

Time-lapse experiments were performed in cell lines following the same initial coating and seeding procedures described for the EdU staining assay. Cells with biosensors were seeded and cultured in Gibco FluoroBrite medium. After allowing cells to attach, drugs were administered as described and maintained throughout live-cell imaging on the Leica Thunder inverted fluorescence microscope (described above) equipped with a cage incubator (OkoLab) to maintain a constant humidified environment at 37°C and 5% CO_2_. Subsequent image processing, tracking, and quantitative analyses are described in the Data analysis section.

### Patient-Derived Organoid culture

Patient-derived organoids (PDOs) were generated in the Institute for Precision Medicine from primary human breast cancer tissue from the Pitt Biospecimen Core consented and collected in accordance with Institutional Review Board protocol STUDY22030183 according to established protocol^87^, with the addition of β-estradiol to the medium. This study used a PDO derived from a primary tumor from a 35-40 female patient with triple negative breast cancer (TNBC) of no special type (NST). Briefly, the tumor was digested with 2 mg/ml collagenase (Sigma) on a rotator, sheared, filtered, and embedded in Cultrex RGF Basement Membrane Extract (R&D Systems) in 24-well Nunc non-treated plates (Fisher), and administered drugs as described. Media was replaced every 2-3 days and PDOs were passaged every 2-4 weeks. Time-lapse imaging experiments were performed as described above to acquire brightfield z-stacks of each PDO condition and maximum intensity projections were generated for each z-stack.

### Confocal imaging

Confocal samples were prepared in 96-well glass-bottom plates following the same coating, seeding, and staining procedures described above. Z-stack imaging was performed using a Nikon AX scanning confocal microscope. For each field of view, optical sections were collected across the full cell volume to generate high-resolution three-dimensional image stacks. Subsequent volumetric z-stack projections were performed as described in the Data Analysis section.

### Colony-forming assay

Cells were seeded in 6-well plates at a density of 1,000–3,000 cells per well. After 24 h, cells were treated with the indicated drugs. At the end of treatment, cells were fixed with 4% PFA) for 30 min, followed by staining with 1% crystal violet solution (Sigma-Aldrich) for 15 min. Plates were washed three times with distilled water and imaged.

### Data Analysis

#### 1. Segmentation, image alignment, and single-cell quantification

Nuclear segmentation was performed on Hoechst images using Cellpose 3.0 (cyto3 model, diameter=None) to generate nuclear masks. Whole-cell masks were obtained by segmenting NaKATPase images, and segmentation quality was inspected using napari^88^. Multiplexed immunofluorescence images were aligned and stitched across imaging rounds using Ashlar^89^. For each cell, nuclear and whole-cell mean intensities were quantified; cytoplasmic intensities were computed by subtracting nuclear from whole-cell signals. Cells were annotated into cell-cycle phases and ploidy states following the criteria described in the Results section. All segmentation, alignment, and quantification scripts are available at: https://github.com/StallaertLab/Thunder_multiplex_pipeline

#### 2. Cell cycle phase and ploidy annotations

Cell cycle phase annotations were assigned to individual cells based on IF measurements according to the following strategy. Gaussian mixture models were used in all cases to distinguish positive and negative expression of biomarkers: (1) using the median nuclear intensity of phospho-RB, cells were labeled as G0 (pRB-) and non-G0 (pRB+). Non-G0 cells were subsequently annotated using median nuclear intensity of phospho-H3 as M (pH3+) and G1/S/G2 (pH3-). Next G1/S/G2 cells were annotated using median nuclear cyclin A2 intensity as G1 (cycA2-) and S/G2 (cycA2+), followed by median nuclear Plk1 intensity as S (PLK1-) and G2 cells (PLK1+). A schematic of this strategy is shown in **Extended Data Figure 1g**. A similar strategy was used to annotate ploidy. First, phospho-RB was used to distinguish G0 from non-G0 as described above. G0 cells were then assigned based on DNA content as diploid (2N) and polyploid (>2N). Non-G0 cells were annotated according to DNA content, with cells with 2N classified as diploid and cells >4N classified as polyploid. Cells with intermediate DNA content (>2N–4N) were further classified using Cyclin A2 expression as diploid (cycA2+) and polyploid (cycA2-). A schematic of this strategy is shown in **Extended Data Figure 41.**

#### 3. Cell state transitions

To quantify the proportion of cells in cell cycle phases of the mitotic and endoreplication cycle, phospho-RB, cyclin A2, and DNA content (Hoeschst) measurements obtained by IF were used to label single cells in: diploid G0 (DNA=2N/pRB-/cycA2-), diploid G1 (DNA=2N/pRB+/cycA2-), diploid S/G2 (DNA=4N/pRB+/cycA2+), polyploid G0 (DNA=4N/pRB-/cycA2-), polyploid G1 (DNA=4N/pRB+/cycA2-), and polyploid S/G2 (DNA=8N/pRB+/cycA2+)

#### 4. Cell cycle mapping

Random-forest feature selection was applied to rank markers contributing to cell cycle and ploidy discrimination as previously described^47^. Selected features were embedded using PHATE^50^ to generate continuous cell cycle maps. Implementation details follow the publicly available pipelines:https://github.com/StallaertLab/multiplex_pipeline, https://github.com/StallaertLab/cc_mapping.

#### 5. Trajectory Inference Analysis

Pseudotime analysis using diffusion pseudotime (DPT) was implemented in Scanpy. We first subset cells to include only diploid G2, diploid M, and polyploid G0 populations. To define the root of the trajectory, we identified the diploid G2 cell closest to the centroid of all diploid S cells in feature space. We then computed diffusion pseudotime, which orders cells along their progression from this root. To resolve the bifurcation between mitotic and polyploid trajectories, we selected terminal cells for each fate manually, as the furthest cell from the root cell along the mitotic vs. polyploid trajectory. We then assigned cells probabilistically to each trajectory branch based on diffusion map distances to the two terminal states, with probabilities normalized by a Gaussian kernel. Branch-specific pseudotime was computed by scaling global DPT values to each terminal state and weighting by branch probability. Trajectory confidence was quantified as 1 minus the Shannon entropy of branch probabilities. Feature dynamics along each trajectory were smoothed using a Gaussian filter applied to cells ordered by pseudotime within each branch.

#### 6. Single cell/organoid tracking and analysis of time-lapse data

For cell line experiments, PCNA-mTq2 images were segmented frame-by-frame using Cellpose 3.0. For PDO experiments, whole organoids were segmented from label-free brightfield images using MobileSAM^90^. For both cells and PDOs, object tracking was performed using Ultrack^91^, followed by manual curation and correction using the Track Gardener plugin for napari. The curated tracks were used to compute CDK2 activity dynamics, annotate cell cycle phase transitions, and measure single-cell cell cycle lengths across the full time-lapse experiment. Cell fate for individual cell traces was annotated manually according to the following criteria: Mitosis: cell division event observed. G0 arrest: cells enter a state with low CDK activity immediately after cell division and remain in this state for duration of experiment. G2 arrest: cells perform a single S phase and then remain in G2 with elevated CDK activity (indicative of an actively proliferating cell) until end of experiment. Endocycling: cells exit from G2 (G2→G0 transition observed). Endomitosis: cells exit from M (M→G0 transition observed). Apoptosis: cells round up and die.

For PDOs, area measurements were extracted and transformed to volume estimations assuming perfect sphericity:

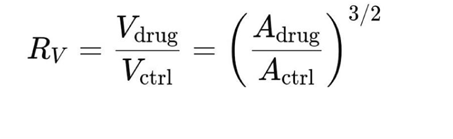

where *R_V_*represents the fold-change in PDO volume for a given timepoint compared to day 0, *V* is volume and *A* is area.

Tracking and annotation tools are available at: https://github.com/fjorka/track_gardener

#### 7. Analysis of colony-forming assay

Total colony area was quantified from crystal violet–stained plates using FIJI (ImageJ). Briefly, images were converted to grayscale, and the stained colony area was segmented using an intensity threshold. The colony-forming capacity was quantified as the ratio of crystal violet–positive area to the total well area.

#### 8. Statistical analysis

All experiments were performed with three biological replicates and statistical significance was evaluated using a paired Student’s *t*-test for two-group comparisons or one-way ANOVA followed by appropriate *post hoc* tests for multiple-group comparisons, unless otherwise stated. Significance: P < 0.05 (*), < 0.01 (**), < 0.001 (***).

## ACKNOWLEDGMENTS

This research was supported by a Research Scholar Grant from the American Cancer Society (RSG-12-345—01-ABC), the Competitive Medical Research Fund of the UPMC Health System, the National Cancer Institute of the National Institutes of Health under award number P30CA047904, by the Chan Zuckerberg Initiative DAF (2023-329680), an advised fund of Silicon Valley Community Foundation, the University of Pittsburgh Organoid Research Core Facility (RRID:SCR_025698), and the UPMC Hillman Cancer Center Cytometry Facility (RRID:SCR_025361). We would like to thank Betsy Ann Varghese and Narayanan Nampoothiri (University of Pittsburgh) for their technical assistance, Dr. Elise Fourquerel (University of Pittsburgh) for assistance in preparing the manuscript, and Drs. Steffi Oesterreich and Adrian Lee (University of Pittsburgh) for their generous gift of the breast cancer cell lines used in this manuscript.

---

**Table S1.**
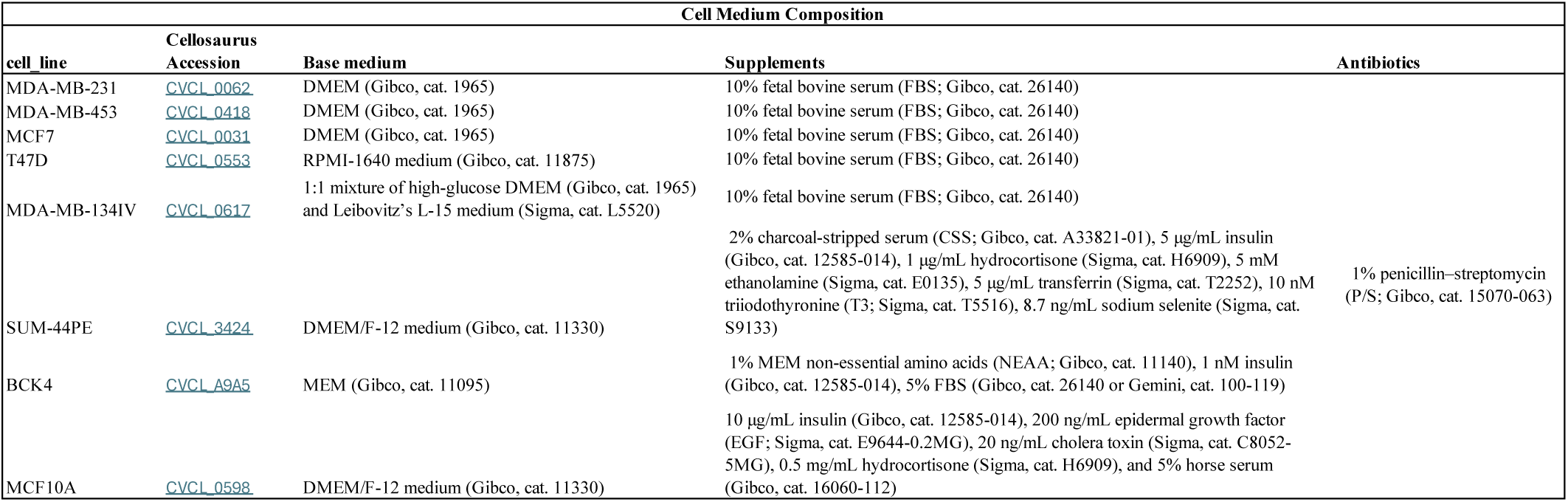

**Table S2.**
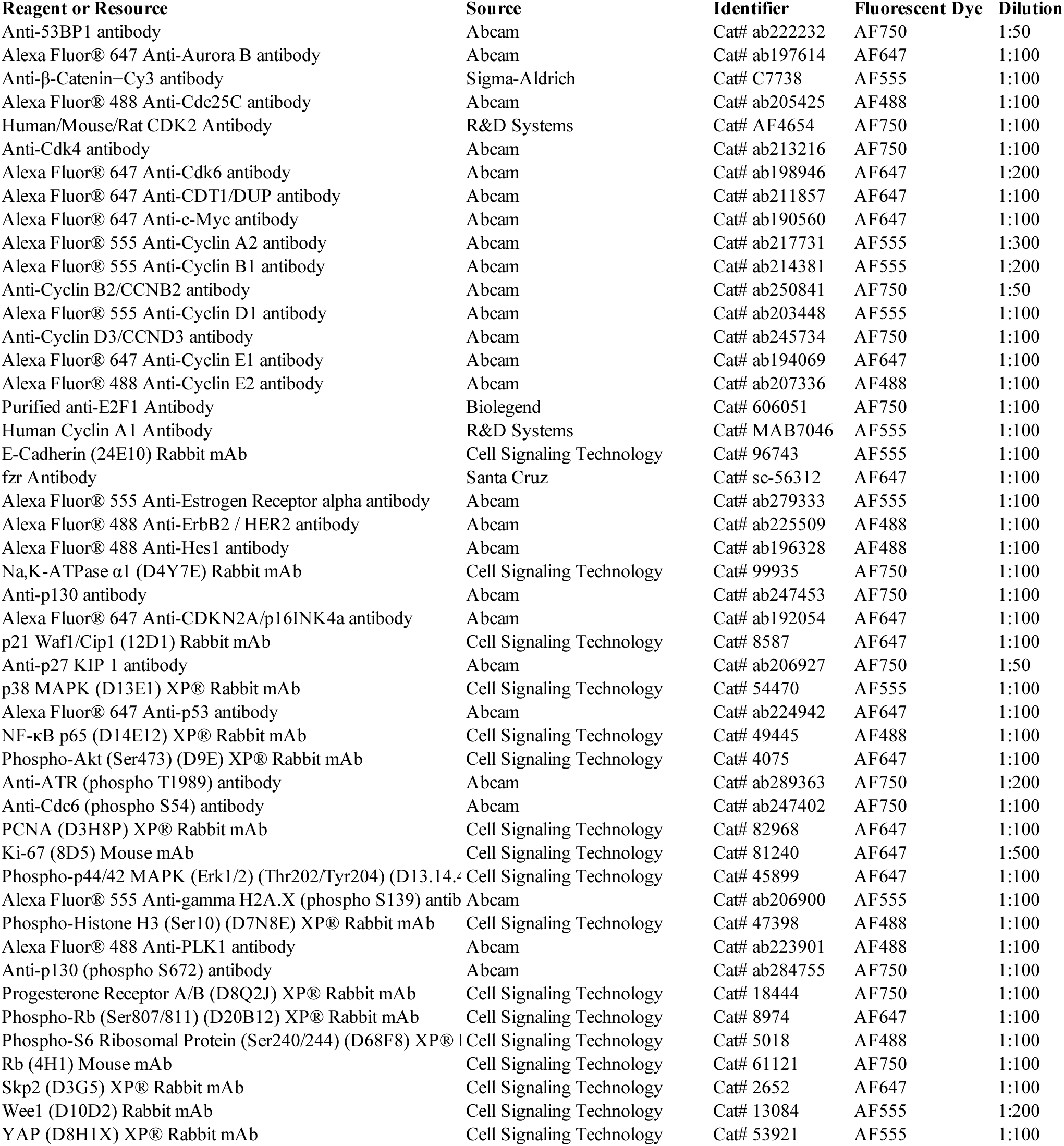

## REFERENCES

1. Ohno, S. Evolution by Gene Duplication. (Springer Nature, 1970).

2. Marlétaz, F. et al. The hagfish genome and the evolution of vertebrates. Nature 627, 811–820 (2024).

3. Comai, L. The advantages and disadvantages of being polyploid. Nat. Rev. Genet. 6, 836–846 (2005).

4. Van de Peer, Y., Mizrachi, E. & Marchal, K. The evolutionary significance of polyploidy. Nat. Rev. Genet. 18, 411–424 (2017).

5. Jiao, Y. et al. Ancestral polyploidy in seed plants and angiosperms. Nature 473, 97–100 (2011).

6. Scott, A. L., Richmond, P. A., Dowell, R. D. & Selmecki, A. M. The influence of polyploidy on the evolution of yeast grown in a sub-optimal carbon source. Mol. Biol. Evol. 34, 2690–2703 (2017).

7. Laetitia Chauve, Emma Bazzani, Clément Verdier, Liam Butler, Martha E. Atimise, Aoibhín McGarry, and Aoife McLysaght. Evidence for increased stress resistance due to polyploidy from synthetic autotetraploid Caenorhabditis elegans. bioRxiv 1–39.

8. Gentric, G., Celton-Morizur, S. & Desdouets, C. Polyploidy and liver proliferation. Clin. Res.Hepatol. Gastroenterol. 36, 29–34 (2012).

9. Miyaoka, Y. et al. Hypertrophy and unconventional cell division of hepatocytes underlie liver regeneration. Curr. Biol. 22, 1166–1175 (2012).

10. Øvrebø, J. I. & Edgar, B. A. Polyploidy in tissue homeostasis and regeneration. Development 145, (2018).

11. Duncan, A. W. et al. The ploidy conveyor of mature hepatocytes as a source of genetic variation. Nature 467, 707–710 (2010).

12. Wilkinson, P. D. et al. Polyploid hepatocytes facilitate adaptation and regeneration to chronic liver injury. Am. J. Pathol. 189, 1241–1255 (2019).

13. Duncan, A. W. Aneuploidy, polyploidy and ploidy reversal in the liver. Semin. Cell Dev. Biol. 24, 347–356 (2013).

14. Ravid, K., Lu, J., Zimmet, J. M. & Jones, M. R. Roads to polyploidy: the megakaryocyte example. J. Cell. Physiol. 190, 7–20 (2002).

15. Nagata, Y., Muro, Y. & Todokoro, K. Thrombopoietin-induced polyploidization of bone marrow megakaryocytes is due to a unique regulatory mechanism in late mitosis. J. Cell Biol. 139, 449–457 (1997).

16. Vainchenker, W. & Raslova, H. Megakaryocyte polyploidization: role in platelet production. Platelets 31, 707–716 (2020).

17. Singh, V. P. et al. Myc promotes polyploidy in murine trophoblast cells and suppresses senescence. Development 150, (2023).

18. Hayakawa, K. et al. Nucleosomes of polyploid trophoblast giant cells mostly consist of histone variants and form a loose chromatin structure. Sci. Rep. 8, 5811 (2018).

19. De Chiara, L., Conte, C., Antonelli, G. & Lazzeri, E. Tubular cell cycle response upon AKI: Revising old and new paradigms to identify novel targets for CKD prevention. Int. J. Mol. Sci. 22, 11093 (2021).

20. De Chiara, L. et al. Tubular cell polyploidy protects from lethal acute kidney injury but promotes consequent chronic kidney disease. Nat. Commun. 13, 5805 (2022).

21. De Chiara, L. & Romagnani, P. Polyploid tubular cells and chronic kidney disease. Kidney Int. 102, 959–961 (2022).

22. Airik, M. et al. Persistent DNA damage underlies tubular cell polyploidization and progression to chronic kidney disease in kidneys deficient in the DNA repair protein FAN1. Kidney Int. 102, 1042–1056 (2022).

23. Patterson, M. et al. Frequency of mononuclear diploid cardiomyocytes underlies natural variation in heart regeneration. Nat. Genet. 49, 1346–1353 (2017).

24. Derks, W. & Bergmann, O. Polyploidy in cardiomyocytes: Roadblock to heart regeneration? Circ. Res. 126, 552–565 (2020).

25. Hirose, K. et al. Evidence for hormonal control of heart regenerative capacity during endothermy acquisition. Science 364, 184–188 (2019).

26. González-Rosa, J. M. et al. Myocardial polyploidization creates a barrier to heart regeneration in zebrafish. Dev. Cell 44, 433–446.e7 (2018).

27. Bielski, C. M. et al. Genome doubling shapes the evolution and prognosis of advanced cancers. Nat. Genet. 50, 1189–1195 (2018).

28. Dewhurst, S. M. et al. Tolerance of whole-genome doubling propagates chromosomal instability and accelerates cancer genome evolution. Cancer Discov. 4, 175–185 (2014).

29. Lv, Y. et al. Differential whole-genome doubling based signatures for improvement on clinical outcomes and drug response in patients with breast cancer. Heliyon 10, e28586 (2024).

30. Zhang, X. et al. Targeting polyploid giant cancer cells potentiates a therapeutic response and overcomes resistance to PARP inhibitors in ovarian cancer. Sci. Adv. 9, eadf7195 (2023).

31. Bojko, A. et al. Improved autophagic flux in escapers from doxorubicin-induced senescence/polyploidy of breast cancer cells. Int. J. Mol. Sci. 21, 6084 (2020).

32. Czarnecka-Herok, J. et al. Therapy-induced senescent/polyploid cancer cells undergo atypical divisions associated with altered expression of meiosis, spermatogenesis and EMT genes. Int. J. Mol. Sci. 23, 8288 (2022).

33. Carroll, C. et al. Drug-resilient Cancer Cell Phenotype Is Acquired via Polyploidization Associated with Early Stress Response Coupled to HIF2α Transcriptional Regulation. Cancer Res. Commun. 4, 691–705 (2024).

34. Prasad, K. & Ben-David, U. A balancing act: how whole-genome doubling and aneuploidy interact in human cancer. Oncotarget 14, 382–383 (2023).

35. Conway, P. J., Dao, J., Kovalskyy, D., Mahadevan, D. & Dray, E. Polyploidy in cancer: Causal mechanisms, cancer-specific consequences, and emerging treatments. Mol. Cancer Ther. 23, 638–647 (2024).

36. Kim, C.-J. et al. Nuclear morphology predicts cell survival to cisplatin chemotherapy. Neoplasia 42, 100906 (2023).

37. Lin, K.-C. et al. The role of heterogeneous environment and docetaxel gradient in the emergence of polyploid, mesenchymal and resistant prostate cancer cells. Clin. Exp. Metastasis 36, 97–108 (2019).

38. Pommier, Y., O’Connor, M. J. & de Bono, J. Laying a trap to kill cancer cells: PARP inhibitors and their mechanisms of action. Sci. Transl. Med. 8, 362ps17–362ps17 (2016).

39. Gopal, A. A., Fernandez, B., Delano, J., Weissleder, R. & Dubach, J. M. PARP trapping is governed by the PARP inhibitor dissociation rate constant. Cell Chem. Biol. 31, 1373–1382.e10 (2024).

40. Petropoulos, M. et al. Transcription-replication conflicts underlie sensitivity to PARP inhibitors. Nature 628, 433–441 (2024).

41. Hopkins, T. A. et al. PARP1 trapping by PARP inhibitors drives cytotoxicity in both cancer cells and healthy bone marrow. Mol. Cancer Res. 17, 409–419 (2019).

42. Shen, Y., Aoyagi-Scharber, M. & Wang, B. Trapping poly(ADP-ribose) polymerase. J. Pharmacol. Exp. Ther. 353, 446–457 (2015).

43. Shu, Z., Row, S. & Deng, W.-M. Endoreplication: The good, the bad, and the ugly. Trends Cell Biol. 28, 465–474 (2018).

44. Edgar, B. A. & Orr-Weaver, T. L. Endoreplication cell cycles: more for less. Cell 105, 297–306 (2001).

45. Lilly, M. A. & Duronio, R. J. New insights into cell cycle control from the Drosophila endocycle. Oncogene 24, 2765–2775 (2005).

46. Fujimaki, K., Jambhekar, A. & Lahav, G. DNA damage checkpoints balance a tradeoff between diploid- and polyploid-derived arrest failures. Cell Rep. 44, 116348 (2025).

47. Stallaert, W. et al. The structure of the human cell cycle. Cell Syst 13, 230–240.e3 (2022).

48. Stallaert, W. et al. The molecular architecture of cell cycle arrest. Mol. Syst. Biol. 18, e11087 (2022).

49. Zikry, T. M. et al. Cell cycle plasticity underlies fractional resistance to palbociclib in ER+/HER2− breast tumor cells. Proceedings of the National Academy of Sciences 121, e2309261121 (2024).

50. Moon, K. R. et al. Visualizing structure and transitions in high-dimensional biological data. Nat. Biotechnol. 37, 1482–1492 (2019).

51. Was, H. et al. Polyploidy formation in cancer cells: How a Trojan horse is born. Semin. Cancer Biol. 81, 24–36 (2022).

52. Zerjatke, T. et al. Quantitative Cell Cycle Analysis Based on an Endogenous All-in-One Reporter for Cell Tracking and Classification. Cell Rep. 19, 1953–1966 (2017).

53. Yang, H. W. et al. Stress-mediated exit to quiescence restricted by increasing persistence in CDK4/6 activation. Elife 9, (2020).

54. Spencer, S. L. et al. The proliferation-quiescence decision is controlled by a bifurcation in CDK2 activity at mitotic exit. Cell 155, 369–383 (2013).

55. Thorn, C. F. et al. Doxorubicin pathways: pharmacodynamics and adverse effects. Pharmacogenet. Genomics 21, 440–446 (2011).

56. Kciuk, M. et al. Doxorubicin-an agent with multiple mechanisms of anticancer activity. Cells 12, 659 (2023).

57. Brito, D. A. & Rieder, C. L. Mitotic checkpoint slippage in humans occurs via cyclin B destruction in the presence of an active checkpoint. Curr. Biol. 16, 1194–1200 (2006).

58. Lok, T. M. et al. Mitotic slippage is determined by p31comet and the weakening of the spindle-assembly checkpoint. Oncogene 39, 2819–2834 (2020).

59. Zeng, J., Hills, S. A., Ozono, E. & Diffley, J. F. X. Cyclin E-induced replicative stress drives p53-dependent whole-genome duplication. Cell 186, 528–542.e14 (2023).

60. el-Deiry, W. S., et al. WAF1, a potential mediator of p53 tumor suppression. Cell 75, 817–825 (1993).

61. Wasielewski, M., Elstrodt, F., Klijn, J. G. M., Berns, E. M. J. J. & Schutte, M. Thirteen new p53 gene mutants identified among 41 human breast cancer cell lines. Breast Cancer Res. Treat. 99, 97–101 (2006).

62. Hayashi, K. et al. Polyploidy mitigates the impact of DNA damage while simultaneously bearing its burden. Cell Death Discov. 10, 436 (2024).

63. Wang, Y., Jha, A. K., Chen, R., Doonan, J. H. & Yang, M. Polyploidy-associated genomic instability in Arabidopsis thaliana. Genesis 48, 254–263 (2010).

64. Herriage, H. C., Hughes, C. L., Fahey, S. K. & Calvi, B. R. Unscheduled polyploidy synergizes with oncogenic mutations to enhance genome instability and tumorigenesis. Cancer Lett. 218008 (2025).

65. Storchova, Z. & Pellman, D. From polyploidy to aneuploidy, genome instability and cancer. Nat. Rev. Mol. Cell Biol. 5, 45–54 (2004).

66. McPherson, A. et al. Ongoing genome doubling shapes evolvability and immunity in ovarian cancer. Nature 644, 1078–1087 (2025).

67. Hobor, S. et al. Mixed responses to targeted therapy driven by chromosomal instability through p53 dysfunction and genome doubling. Nat. Commun. 15, 4871 (2024).

68. Gemble, S. et al. Genetic instability from a single S phase after whole-genome duplication. Nature (2022) doi:10.1038/s41586-022-04578-4.

69. Utani, K.-I., Okamoto, A. & Shimizu, N. Generation of micronuclei during interphase by coupling between cytoplasmic membrane blebbing and nuclear budding. PLoS One 6, e27233 (2011).

70. Panagaki, D. et al. Nuclear envelope budding is a response to cellular stress. Proc. Natl. Acad. Sci. U. S. A. 118, e2020997118 (2021).

71. Keuenhof, K. S. et al. Nuclear envelope budding and its cellular functions. Nucleus 14, 2178184 (2023).

72. Crasta, K. et al. DNA breaks and chromosome pulverization from errors in mitosis. Nature 482, 53–58 (2012).

73. Sundaram, M., Guernsey, D. L., Rajaraman, M. M. & Rajaraman, R. Neosis: a novel type of cell division in cancer. Cancer Biol. Ther. 3, 207–218 (2004).

74. Zhang, Z. et al. Irradiation-induced polyploid giant cancer cells are involved in tumor cell repopulation via neosis. Mol. Oncol. 15, 2219–2234 (2021).

75. Kwon, M., Leibowitz, M. L. & Lee, J.-H. Small but mighty: the causes and consequences of micronucleus rupture. Exp. Mol. Med. 52, 1777–1786 (2020).

76. Bukkuri, A. et al. A life history model of the ecological and evolutionary dynamics of polyaneuploid cancer cells. Sci. Rep. 12, 13713 (2022).

77. Bukkuri, A. et al. Stochastic models of Mendelian and reverse transcriptional inheritance in state-structured cancer populations. Sci. Rep. 12, 13079 (2022).

78. Lee, H. O., Davidson, J. M. & Duronio, R. J. Endoreplication: polyploidy with purpose. Genes Dev. 23, 2461–2477 (2009).

79. Fox, D. T. & Duronio, R. J. Endoreplication and polyploidy: insights into development and disease. Development 140, 3–12 (2013).

80. Guerrero Zuniga, A., et al. Sustained ERK signaling promotes G2 cell cycle exit and primes cells for whole-genome duplication. Dev. Cell 59, 1724–1736.e4 (2024).

81. Kim, G.-Y. et al. The stress-activated protein kinases p38 alpha and JNK1 stabilize p21(Cip1) by phosphorylation. J. Biol. Chem. 277, 29792–29802 (2002).

82. McKenney, C. et al. CDK4/6 activity is required during G_2 arrest to prevent stress-induced endoreplication. Science 384, eadi2421 (2024).

83. Iyengar, M. et al. CDK4/6 inhibition as maintenance and combination therapy for high grade serous ovarian cancer. Oncotarget 9, 15658–15672 (2018).

84. Coffman, L. G., et al. Phase I trial of ribociclib with platinum chemotherapy in ovarian cancer. JCI Insight 7, (2022).

85. Decker, T. et al. Anti-hormonal maintenance treatment with the CDK4/6 inhibitor ribociclib after 1st line chemotherapy in hormone receptor positive / HER2 negative metastatic breast cancer: A phase II trial (AMICA). Breast 72, 103575 (2023).

86. Stringer, C. & Pachitariu, M. Cellpose3: one-click image restoration for improved cellular segmentation. Nat. Methods 22, 592–599 (2025).

87. Sachs, N. et al. A living biobank of breast cancer organoids captures disease heterogeneity. Cell 172, 373–386.e10 (2018).

88. Chiu, C.-L., Clack, N. & the napari community. Napari: A python multi-dimensional image viewer platform for the research community. Microsc. Microanal. 28, 1576–1577 (2022).

89. Muhlich, J. L. et al. Stitching and registering highly multiplexed whole-slide images of tissues and tumors using ASHLAR. Bioinformatics 38, 4613–4621 (2022).

90. Zhang, C., et al. Faster segment anything: Towards lightweight SAM for mobile applications. arXiv [cs.CV] (2023).

91. Bragantini, J. et al. Ultrack: pushing the limits of cell tracking across biological scales. Nat. Methods 22, 2423–2436 (2025).

